# Structure and Function of Gli123 Involved in *Mycoplasma mobile* Gliding

**DOI:** 10.1101/2022.09.07.507039

**Authors:** Daiki Matsuike, Yuhei O Tahara, Takahiro Nonaka, Heng Ning Wu, Tasuku Hamaguchi, Hisashi Kudo, Yuuki Hayashi, Munehito Arai, Makoto Miyata

**Affiliations:** Graduate School of Science, Osaka Metropolitan University, Sumiyoshi-ku, Osaka, Japan; The OCU Advanced Research Institute for Natural Science and Technology (OCARINA), Osaka Metropolitan University, Sumiyoshi-ku, Osaka, Japan; Graduate School of Science, Osaka City University, Sumiyoshi-ku, Osaka, Japan; Institute of Multidisciplinary Research for Advanced Materials (IMRAM), Tohoku University, Aoba-ku, Miyagi, Japan; Department of Life Sciences, The University of Tokyo, Meguro Tokyo, Japan; Graduate School of Science, Technology and Innovation, Kobe University, Nada, Kobe, Japan; Environmental Science Center, The University of Tokyo, Bunkyo, Tokyo, Japan; Department of Physics, The University of Tokyo, Meguro, Tokyo, Japan

**Keywords:** electron microscopy, small-angle X-ray scattering, repeat sequence, single-particle analysis, mollicutes, lipoprotein, conformational change, motility

## Abstract

*Mycoplasma mobile* is a fish pathogen that glides on solid surfaces by means of a unique mechanism. The gliding machinery of *M. mobile* is composed of internal and surface structures. In the present study, we focused on the function and structure of Gli123, a surface protein that is essential for the localization of other surface proteins. The amino acid sequence of Gli123, which is 1128 amino acids long, contains lipoprotein-specific repeats. We isolated the native Gli123 protein from *M. mobile* cells and a recombinant protein, rGli123, from *Escherichia coli*. The isolated rGli123 complemented a non-binding and non-gliding mutant of *M. mobile* that lacked Gli123. Circular dichroism and rotary-shadowing electron microscopy (EM) showed that rGli123 has a structure that is not significantly different from that of the native protein. Rotary-shadowing EM suggested that the molecules changed their shape between globular and rod-like structures, depending on the ionic strength of the solution. Negative-staining EM coupled with single-particle analysis revealed that Gli123 forms a globular structure featuring a small protrusion with dimensions of 20.0, 14.5, and 16.0 nm. Small-angle X-ray scattering analyses indicated a rod-like structure composed of several tandem globular domains with total dimensions of approximately 34 nm length and 4 nm width. Both molecular structures were suggested to be dimers based on the predicted molecular size and structure. Gli123 may have evolved by multiplication of repeating lipoprotein units and acquired clumping role of surface proteins.

**IMPORTANCE:** *Mycoplasmas* are pathogenic bacteria that are widespread in animals. They are characterized by small cell and genome sizes but are equipped with unique abilities to escape host immunity, such as surface variation and gliding. Here, we focused on a surface-localizing protein that is essential for *Mycoplasma mobile* gliding. The findings of this study suggested that the protein undergoes drastic conformational changes between its rod-like and globular structures. These changes may be caused by a repetitive structure common in the surface proteins that is responsible for the modulation of the cell surface structure and related to the assembly process for the surface gliding machinery. An evolutionary process for this unique mycoplasma gliding mechanism has also been suggested in the present study.

## INTRODUCTION

Class Mollicutes, which includes the genus *Mycoplasma*, consists of organisms fit to parasitic or commensal life cycles, as represented by their small genome size, lack of peptidoglycan layer, and antigenic modulation (1–3). Interestingly, more than 20 *Mycoplasma* species are known to glide on solid surfaces through unique mechanisms (4). *Mycoplasma* gliding mechanisms can be classified into two types, as represented by either *Mycoplasma mobile* or *Mycoplasma pneumoniae* (4, 5). Four species are known to perform mobile-type gliding, including *M. mobile, Mycoplasma pulmonis, Mycoplasma testudineum*, and *Mycoplasma agassizii*, which serve as pathogens for freshwater fish, mice, turtles, and turtles, respectively (6, 7). The gliding mechanism has been studied with a focus on *M. mobile*, which glides at speeds of 2.0 to 4.5 μm/s (4, 8). The gliding machinery, which forms as a protrusion, can be divided into two parts: the inside and surface structures (4, 9, 10). Its surface structure is composed of four proteins: Gli123, Gli349, Gli521, and Gli42. Gli123, Gli349, and Gli521 have been reported as being essential for binding and gliding, based on analyses of mutants missing each of these proteins (11–13). Gli349 is a flexible rod-shaped, 95-nm-long protein with a C-terminal globular domain (11, 14, 15). It catches sialylated oligosaccharides on the surfaces of hosts as a “leg” for gliding, and pulls the cell (16, 17). Gli521 is a rather rigid rod-shaped 120-nm-long protein with N-terminal globular and C-terminal hook domains (18, 19) that is suggested to transmit the gliding force generated in an internal motor to Gli349, like a “crank” (5, 18, 19). Gli123 localizes Gli349 and Gli521 correctly to the gliding machinery, as a “mount” (13). Gli349 and Gli521 are conserved in all four species of *M. mobile, M. pulmonis, M. testudineum*, and *M. agassizii*, whereas Gli123 and Gli42 are only found in *M. mobile* and *M. pulmonis* (20). In this study, we isolated Gli123 and found that a mutant lacking this protein could be complemented by the isolated protein instead. We analyzed Gli123 structures using electron microscopy (EM), small-angle X-ray scattering (SAXS), and other biochemical methods, and suggested a plausible structural model and its conversion, which may be related to the role of the protein in gliding motility.

## RESULTS

### Gli123 orthologs in the *Mycoplasma* species

Orthologs of *gli123* and *gli42* genes were not found in *M. agassizii* and *M. testudineum*, unlike the orthologs of nine other genes involved in *M. mobile* gliding (20). The orthologs of *gli349* and *gli521* from *M. testudineum* and *M. agassizii* have been found to be positioned closer to *M. pulmonis* than to *M. mobile* in the phylogenetic tree (20). Therefore, we performed a PSI-BLAST search (21) using the amino acid sequences of Gli123 and Gli42 of *M. pulmonis*, and found the orthologs of *gli123* and *gli42* in the genomes of *M. agassizii* and *M. testudineum* (Table S1). The phylogenetic trees constructed for *gli123, gli42, gli349*, and *gli521* genes using the maximum likelihood method shared their topology, as did the phylogenetic tree of 16S ribosomal DNA sequences (Fig. S1) (20). Next, we examined the localization of these genes in the genome. In *M. mobile* and *M. pulmonis*, the genes were clustered in one locus (13, 20), while in *M. agassizii* and *M. testudineum*, they were split into two loci, *gli123-gli42* and *gli349-gli521*, although the actual distances are unknown in the unassembled genome sequences (Fig. S2).

### Sequence analysis of Gli123 and the lipoprotein-17 domain

In this study, we found that the amino acid sequences of Gli123 protein and its orthologs include four to six lipoprotein-17 domains, which are composed of approximately 100 amino acids (Fig. 1A) (22, 23). The lipoprotein-17 domain is found in 233 proteins of 43 bacterial species, mostly in lipoproteins that play a role in antigenic variation or adhesion. In *Mycoplasma*, the lipoprotein-17 domain was identified in 221 proteins from 32 species (Table S2). This domain was found in 15 Mvsps, Gli123, and Gli349 in *M. mobile* (23, 24). The Gli123 protein of *M. mobile* had four lipoprotein-17 domains, while that of *M. pulmonis* and *M. testudineum* had five and that of *M. agassizii* had six (Fig. 1A and Table S3). Mvsps have a number of lipoprotein-17 domains, up to 16 (Fig. 1A and Table S3). This may suggest that the surface proteins adjust their lengths by means of repetitive fusion of the domain (4). The hidden Markov model scores for the lipoprotein-17 domain were distributed between 30 and 70 for the Mvsps repeats and between 10 and 30 for the Gli123 and Gli349 repeats (Fig. 1B).

**FIG. 1.**
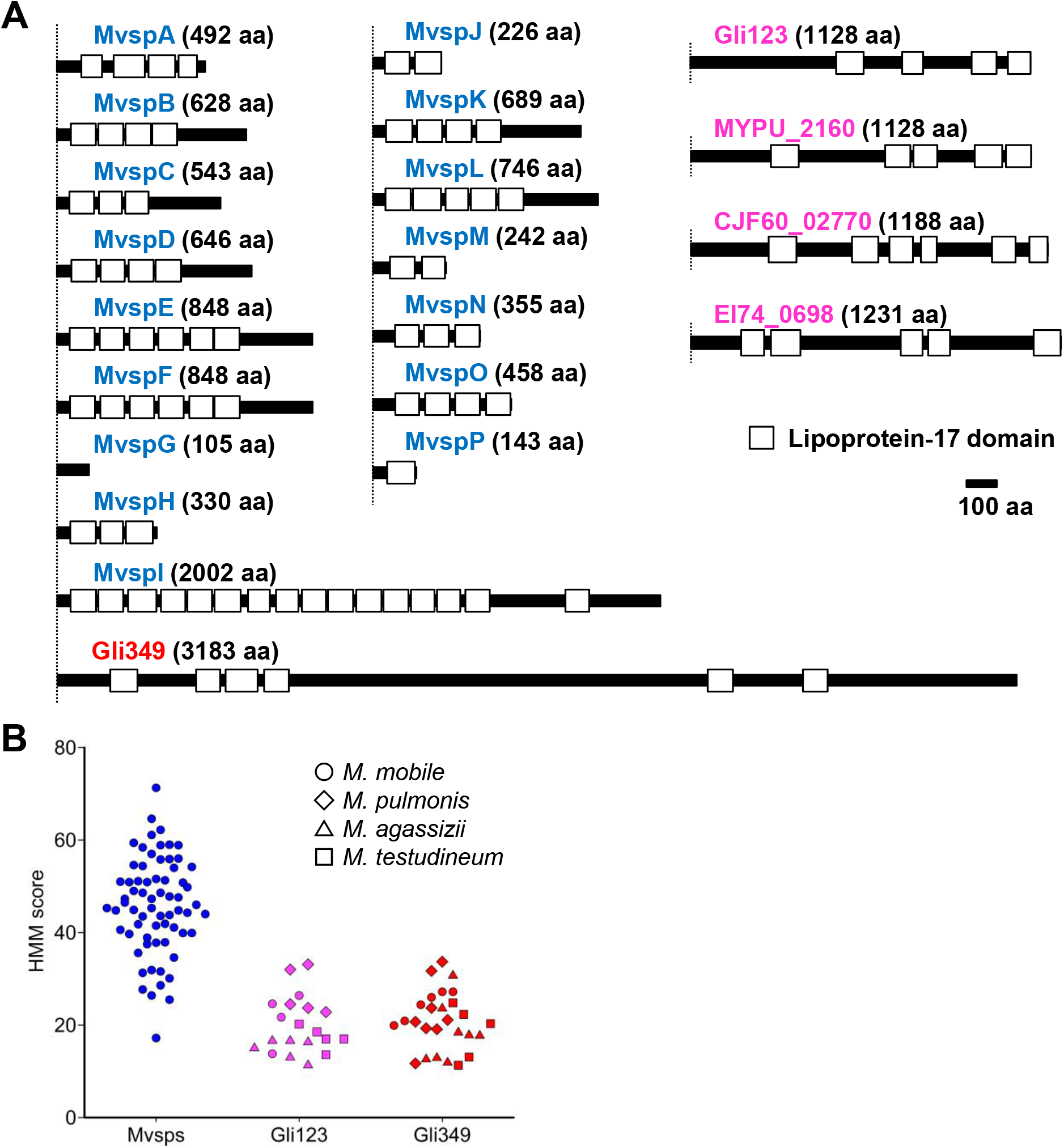
Lipoprotein-17 domains in Mvsps, Gli123, and Gli349. (A) Distributions of lipoprotein-17 domains in Mvsps, Gli349, and Gli123 sequences have been shown using white boxes. MYPU_2160, CJF60_02770, and EI74_0698 are Gli123 orthologs of *M. pulmonis*, *M. agassizii*, and *M. testudineum*, respectively. (B) Column distribution of the scores calculated using the hidden Markov model (HMM) for each lipoprotein-17 domain. The scores for each lipoprotein-17 domain of Gli123 and Gli349 and orthologs in *M. mobile, M. pulmonis, M. agassizii*, and *M. testudineum* and Mvsps in *M. mobile* are shown.

### MvspI mutant strains of *M. mobile*

The isolation of native Gli123 (nGli123) from *M. mobile* cell lysates is a long process because the lipoprotein, MvspI encoded as MMOB3340, behaves in a similar way in the isolation process. MvspI is characterized by a mass of 220 kDa, with 16 copies of lipoprotein-17 domains (24). We isolated a spontaneous MvspI-deficient mutant to simplify the isolation process of Gli123. Colonies that were non-reactive to a monoclonal antibody against MvspI were screened and isolated through colony blotting and Amido Black staining (Fig. S3A). Two colonies that were non-reactive to the antibody were isolated from an approximate of 10000 colonies. Sodium dodecyl sulfate-polyacrylamide gel electrophoresis (SDS-PAGE) analysis of the isolate revealed a lack of MvspI protein (Fig. S3B). The band intensities of the surface protein, MvspF encoded as MMOB3300, was higher in the isolated mutant than in the wild-type (WT) strain (Fig. S3B). Genome analysis revealed that the *mvspI* gene of the isolate had a frameshift mutation, resulting in a truncated peptide of 97 amino acids (Table S4). In addition to mutations in the *mvspI* gene, nine other mutations were identified in *mvsps* and others (Table S4). This mutant grew constantly through serial cultivation of large volumes, without recovery of MvspI. The mutant cells glided normally, but the culture tended to form clumps, suggesting different surface interactions from those of the WT. This mutant was named the MvspI^-^ strain and used in subsequent experiments.

### Isolation of Gli123 from *M. mobile*

We isolated the nGli123 protein from cultured cells of the *M. mobile* MvspI^-^ strain (Fig. 2A). The collected cells were suspended in a buffer containing 1% Triton™ X-100 and treated on ice, for 1 h. Following that, the lysates were separated by means of centrifugation into a precipitate and a soluble fraction containing nGli123, Gli349, Gli521, and MMOB1650 (Fig. 2B). The soluble fraction was then subjected to anion-exchange chromatography. The nGli123 protein was bound to an anion-exchange column with 20 mM NaCl and eluted using a gradient of NaCl. Gli521 and MMOB1650 were separated from nGli123 by means of this elution process (Fig. 2B). Next, we subjected the fractions containing nGli123 and Gli349 to size-exclusion chromatography (SEC). The proteins, which were traced by means of absorbance, were divided into four peaks, other than the void-volume fraction (Fig. 2C). SDS-PAGE and mass spectrometry identified nGli123 in the second peak near the position of the 150 mL elution (Fig. 2B and 2C). The Stokes radius of nGli123 was calculated as 5.4 nm from the elution position (Fig. 2C). The final yield of the nGli123 protein was 0.3 mg, isolated from 3 L of growth medium. The N-terminal amino acid sequence of the nGli123 protein band collected from the electrophoresis of the whole-cell lysate was analyzed using Edman degradation and determined as Ala-Ile-Ala-Ile-Gly, thereby showing that the N-terminus of 18 amino acids including the partial transmembrane segment of 11 amino acids is truncated in the mature protein (Fig. 2A) (13).

**FIG. 2.**
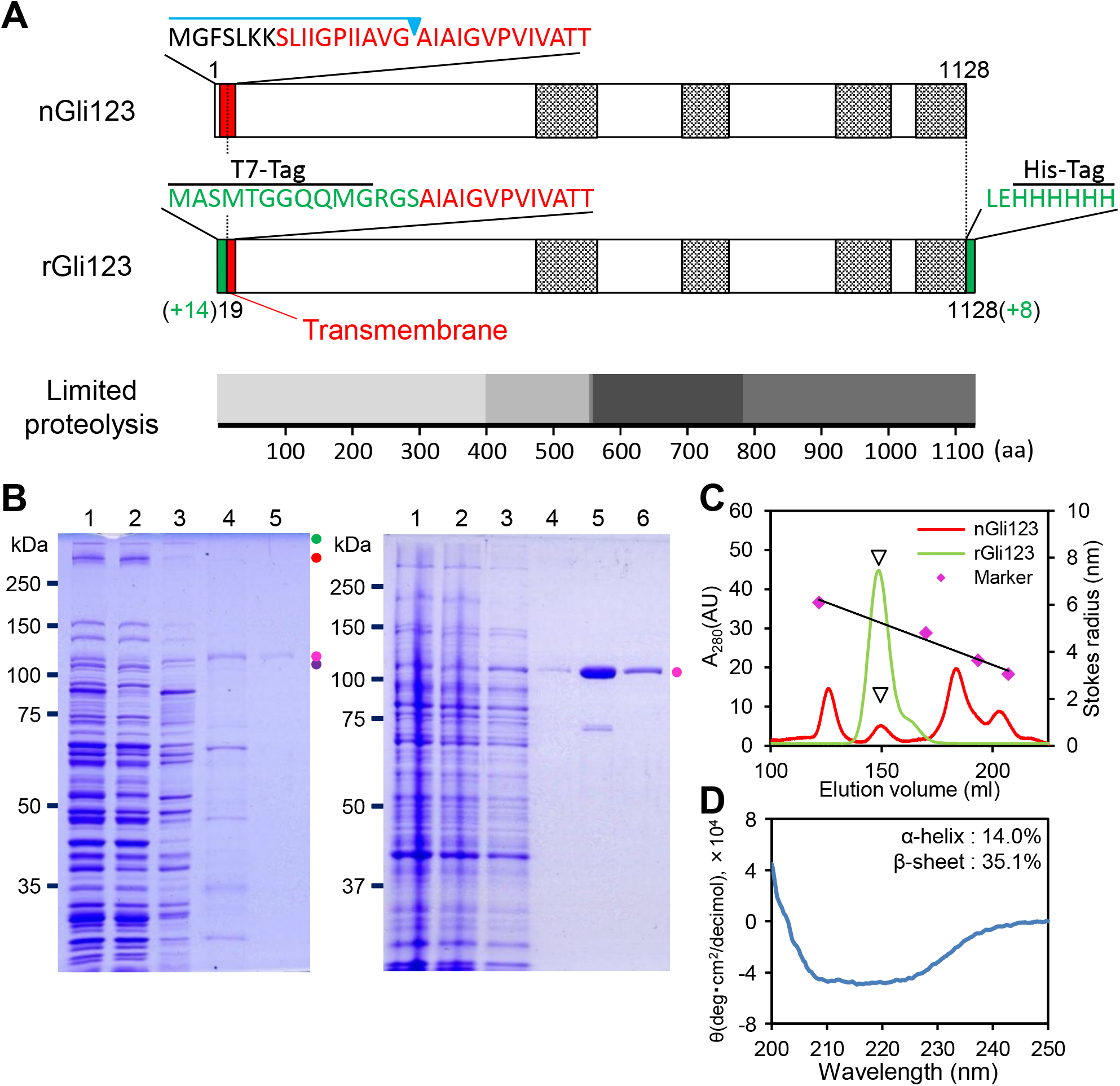
Amino acid sequence and isolation process of native and recombinant Gli123 proteins (nGli123 and rGli123, respectively). (A) Schematic illustration of the amino acid sequences of nGli123 and rGli123. Transmembrane segment and tagging sequences are marked in red and green, respectively. The blue line and triangle indicate the truncated part of the mature protein. The hatched boxes indicate the lipoprotein-17 domains. The bottom diagram for limited proteolysis shows overlap of degradation products in the digest, suggesting rigid areas, as indicated by the four shades. (B) Left: protein profiles of the fractions obtained in the process of isolation of nGli123 from *M. mobile*. Lane 1, whole-cell lysate; Lane 2, Triton™-insoluble fraction; Lane 3, Triton™-soluble fraction; Lane 4, nGli123 fraction eluted by means of anion-exchange chromatography; Lane 5, nGli123 fraction eluted by means of size-exclusion chromatography (SEC). The protein bands of Gli521, Gli349, MMOB1650, and nGli123 are marked using green, red, purple, and magenta circles, respectively. Right: protein profiles of fractions in the rGli123 purification process. Lane 1, whole-cell lysate; Lanes 2 and 3, insoluble and soluble fractions of the lysate after sonication, respectively; Lane 4, rGli123 fraction after Ni^2+^-NTA affinity chromatography; Lanes 5, rGli123 fraction after hydrophobic interaction chromatography; Lanes 6, rGli123 fraction after SEC. The protein fractions were applied to SDS-PAGE with a 10% acrylamide gel and stained with Coomassie Brilliant Blue. The molecular masses are indicated on the left. (C) SEC of nGli123 and rGli123. Thyroglobulin (669 kDa, 8.5 nm), ferritin (440 kDa, 6.1 nm), aldolase (158 kDa, 4.8 nm), and ovalbumin (43 kDa, 3.1 nm) were used as the markers for molecular mass and Stokes radius. The triangles mark the peaks of nGli123 and rGli123. (D) CD spectrum of rGli123.

### Expression and purification of recombinant Gli123

The yield of nGli123 was not high enough to perform various experiments. Thus, we expressed recombinant Gli123 (rGli123) with a 6×His-tag at the C-terminus of mature Gli123 in *E. coli* (Fig. 2A) (13) and purified it using three steps: (i) nickel-nitrilotriacetic acid (Ni^2+^-NTA) affinity chromatography, (ii) hydrophobic interaction chromatography, and (iii) SEC (Fig. 2B). The purified fraction was recovered by means of SEC, as a single peak that included only the rGli123 protein, which was then confirmed using SDS-PAGE and visualized using Coomassie Brilliant Blue (CBB) staining and mass spectrometry (Fig. 2B and 2C). The Stokes radius of rGli123 was estimated as 5.4 nm, from the elution position of SEC (Fig. 2C), which agreed with that for the nGli123 isolated from *M. mobile* cells. The final yield of the rGli123 protein was approximately 10 mg per 1 L of *E. coli* culture. The circular dichroism spectrum of rGli123 showed that the secondary structural contents included 14.0% α-helices and 35.1% β-sheets (Fig. 2D) (25, 26), consistent with the predictions made from the amino acid sequence using XtalPred, *i.e*., 14.0% for α-helices and 43.0% for β-sheets. The purified rGli123 protein was subjected to limited proteolysis with trypsin, to identify the flexible and rigid parts (Fig. 2A). The SDS-PAGE profile of the cleaved fragment showed that the rGli123 protein finally converged into three products through several fragmentation steps (Fig. S4A). Mass spectrometry results showed that the three final fragments (f, g, and h) corresponded at least to the 405–1128 (+8), 565–1128 (+8), and 559–789 amino acid sequences, respectively. Based on these results, we believe that some components of the molecule are rigid, as presented in Fig. 2A.

### Complementation of the binding and gliding abilities of the *M. mobile* mutant with additional Gli123 proteins

To examine the complementation activity of Gli123 for the Gli123-lacking mutant, m12 (13), we added purified rGli123 protein to the cell suspension and incubated it for 3 h at 25°C. The cells were inserted into the tunnel slides with video recordings of phase-contrast optical microscopy (27). m12 cells were unable to bind to the glass surface (13), whereas cells with rGli123 protein showed binding to the glass surface (Fig. 3A). The average number of bound cells in a 100 μm^2^ field on glass was 15.1 ± 3.4 cells *(n* = 15), which was 69.1% of that of the WT. 14.9 ± 8.5% of the bound cells showed unidirectional gliding, similar to the WT strain, with a speed of 1.5 ± 0.8 μm/s (*n* = 45), comparable with that of the WT strain, 2.7 ± 0.5 μm/s (*n* = 45) (Fig. 3B and 3C), suggesting that the rGli123 protein binds and complements the gliding machinery (13). Next, we examined the binding of the rGli123 proteins to the mutant cells. Mutant cells incubated with rGli123 were recovered by means of centrifugation and analyzed using SDS-PAGE (Fig. 3D). This result showed that rGli123 bound to the mutant cells at an amount comparable to that of the WT strain.

**FIG. 3.**
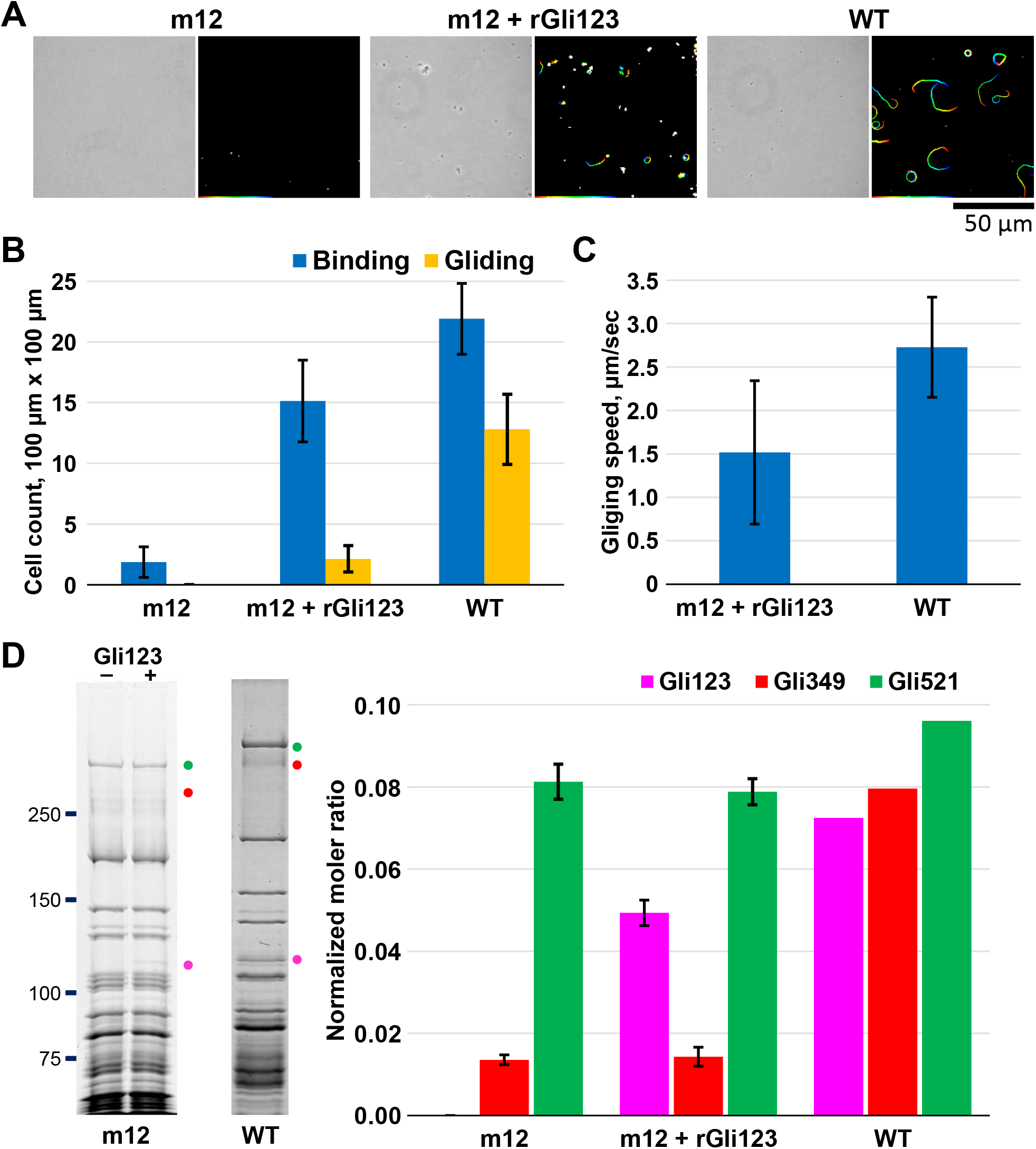
Recovery of gliding and binding by Gli123 proteins in m12 mutant cells. (A) Cell movements analyzed using optical microscopy. The cells were observed using phase-contrast microscopy (left) and a video was recorded. The video frame (at every 0.03 s) was overlaid as a stack for 10 s, changing its color from red to blue (right). (B) Number of cells binding and gliding on glass surfaces. (C) Gliding speeds. (D) Left: Protein profiles of m12 mutant and m12 wild-type cells, after addition of rGli123 protein. The protein bands for Gli521, Gli349, and Gli123 are marked using green, red, and magenta circles, respectively. Right: Molecular ratios of Gli123, Gli349, and Gli521 were calculated from the band intensities normalized to the value of MvspI.

### Conformational changes in Gli123

We observed the structures of the nGli123 and rGli123 proteins using rotary-shadowing EM. In rotary-shadowing EM, protein molecules sprayed on a rotating mica surface are shadowed by platinum particles from a low angle, to observe their molecular shape (19, 24). We observed particles of similar sizes and shapes in the field of view. The nGli123 and rGli123 proteins showed similar rod-like structures, with lengths of 35.6 ± 5.0 and 35.9 ± 4.1 nm, respectively, under conditions including 100 mM ammonium acetate (Fig. 4A and 4B). Next, we observed the nGli123 and rGli123 proteins in the presence of 500 mM ammonium acetate. Interestingly, both proteins showed globular structures with long axes of 21.0 ± 3.2 and 20.1 ± 4.1 nm (Fig. 4A and 4B). We then examined the molecular ratios of the two structures in 100, 250, and 500 mM ammonium acetate. The proportions of rod-like structures at each concentration were 89.7%, 47.0%, and 5.7%, respectively, and the ratio changed from rod-like to globular structures depending on the ammonium acetate concentration (Fig. 4C). Furthermore, we measured the light scattering of rGli123. Light scattering intensity measured at different concentrations increased with ammonium acetate concentration, in the range of 50–650 mM, suggesting that the conformation of rGli123 molecules depends on the concentration of ammonium acetate (Fig. 4D). The 50% effective concentration (EC50) was 261.0 ± 11.3, 238.3 ± 16.1, and 292.9 ± 4.9 mM at protein concentrations of 0.23, 0.75, and 1.23 mg/mL, respectively (Fig. 4D). The changes in the light scattering intensities of rGli123 proteins due to changes in ionic strength were in good agreement with the ratios of globular and rod-like structures measured using rotary-shadowing EM (Fig. 4C and 4D).

**FIG. 4.**
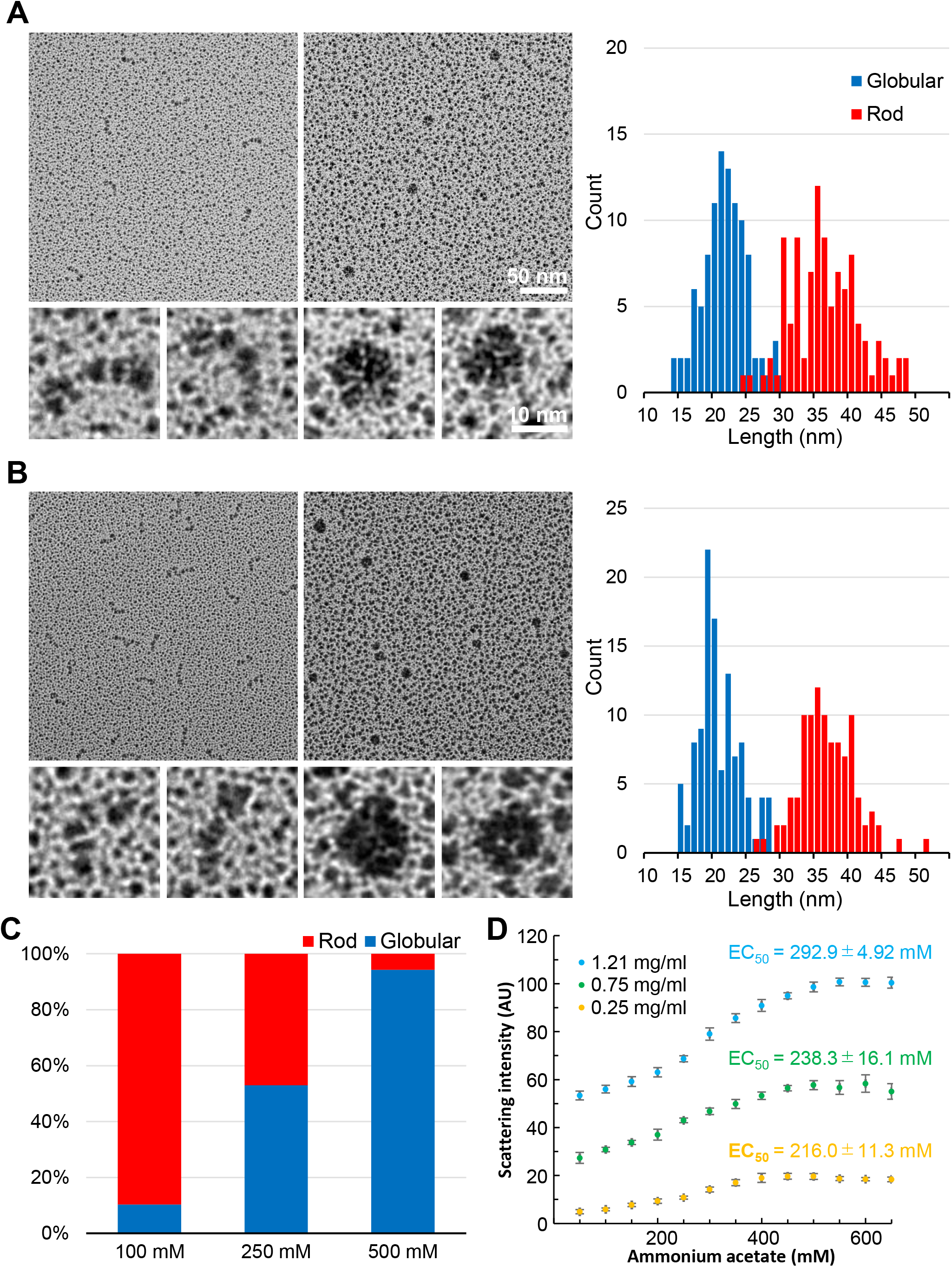
Ionic strength-induced conformational changes in the nGli123 and rGli123 proteins. Field images of the nGli123 (A) and rGli123 (B) proteins under the conditions of 100 mM (left) and 500 mM (right) ammonium acetate. Scale bar: 50 nm. The representative particle images are magnified in the bottom. Scale bar: 10 nm. The distributions of diameter and length are shown for the globular and rod-like structures on the right. (C) Ratio of globular and rod-like structures of the rGli123 protein under the conditions of 100, 250, and 500 mM ammonium acetate. (D) Light scattering of rGli123 proteins, as a function of ammonium acetate concentration.

### Globular structure of Gli123

Negative-staining EM was performed to visualize the three-dimensional structure of globular nGli123 with better resolution. The field images of EM showed uniform globular particles of 15–20 nm (Fig. 5A), and their density changed depending on the protein concentration used. In addition, many rod-like structures were observed in the background. We selected globular particles (Fig. 5A) and obtained 2D-averaged images using the structural analysis software EMAN2.91 (Fig. 5B). The 2D-averaged images showed that the Gli123 molecules had dimensions of approximately 17 and 14 nm for the major and minor axes, respectively. Based on the averaged image of the Gli123 protein, a three-dimensional image was reconstructed, without C1 and with C2, the two-rotational symmetry enforcement. The three-dimensional (3D)-reconstructed globular structure of Gli123 had a protrusion that was reminiscent of a ‘spinning top’ (Fig. 5C). Considering the globular molecular shape, the molecular weight of Gli123 was estimated to be 244 kDa from the elution position of SEC, indicating that the protein behaves as a dimer (Fig. 2C).

**FIG. 5.**
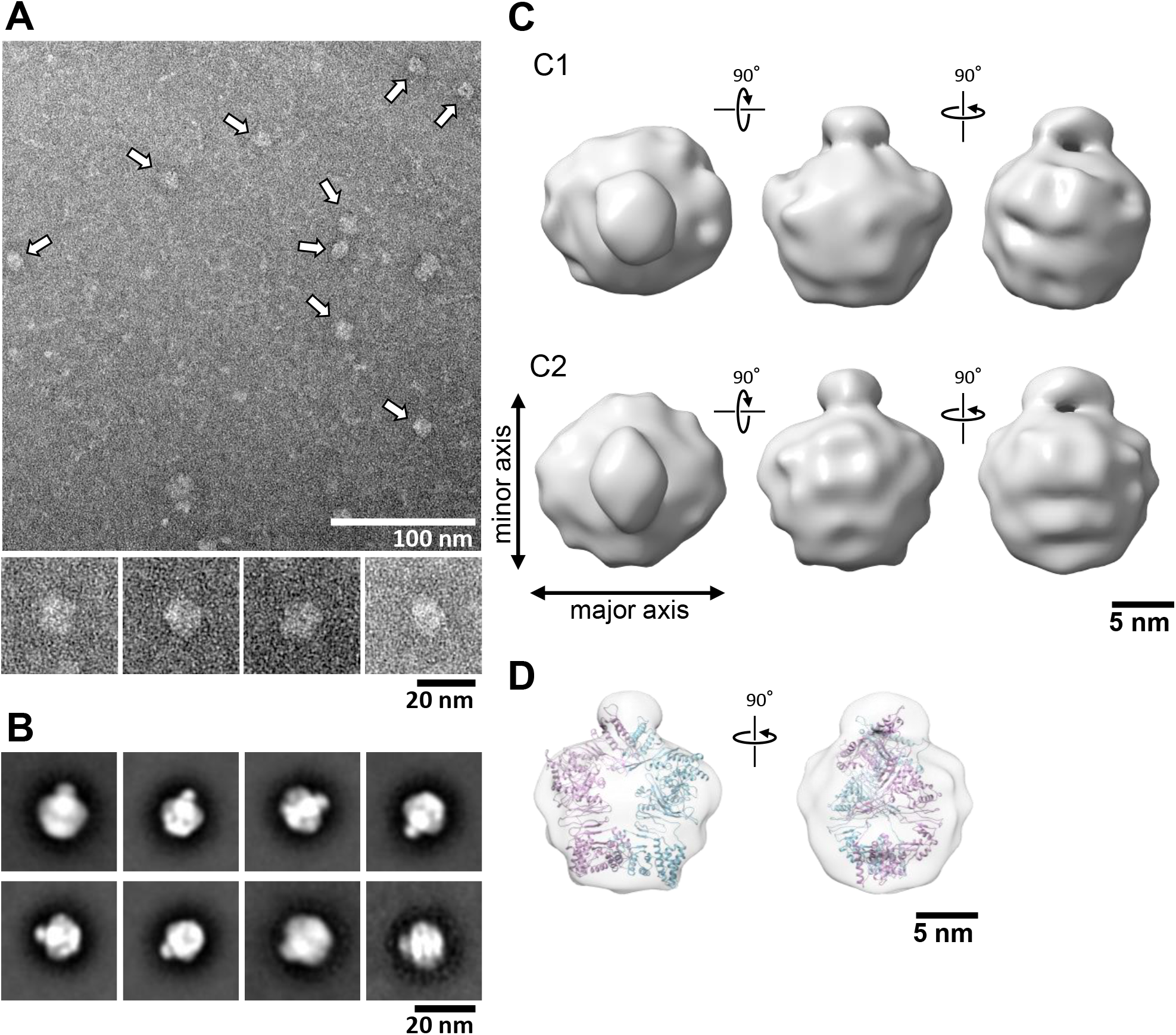
Globular structure of Gli123 protein, as observed by means of negative-staining EM. (A) Field image of Gli123 molecules that were negatively stained with 2% uranyl acetate. Representative particle images from the fields are shown in the bottom. Single-particle analysis was performed using EMAN2.91 software. (B) Representative averaged images of Gli123 were selected from 50 classes which were obtained by averaging approximately 10,000 particles using the command “2D Analysis”. (C) Three-dimensional reconstruction of Gli123 molecule by means of *ab initio* reconstruction with the parameters; C1, non-symmetrical; C2, two-rotational symmetry. (D) Superposition of the C2 model structures obtained from the single-particle analysis and those predicted from the amino acid sequence (1–1000 aa) using Robetta.

### Rod-like structure of Gli123

SAXS was used to obtain the 3D rod-like structure of Gli123. The rGli123 protein under low ionic strength condition was subjected to SEC-SAXS. The protein showed an elution curve with a single peak (Fig. 6A). We analyzed the scattering curve at the peak top of the elution profile with a Guinier plot [ln *I*(*Q*) *vs. Q*^2^, where *I*(*Q*) is the scattering intensity at scattering vector *Q*], to estimate the size of the rGli123 protein (Fig. 6B). The Guinier plot was based on the scattering curve at small angles, and the square of the radius of gyration (*R_g_*) was obtained from the slope of the approximate line in the Guinier region (*Q*×*R*_g_ < 1.3) as ln *I*(*Q*) = ln *I*(0) – *R*_g_^2^*Q*^2^/3, where *I*(0) is the zero-angle scattering intensity (28). *R*_g_ was calculated as 8.24 nm from the slope of the approximate line (Fig. 6B). Next, to discuss the shape of the rGli123 protein, the scattering curve was analyzed using a Kratky plot [*I*(*Q*)*Q*^2^ *vs. Q*] (Fig. 6C), which shows the characteristics of the protein shape (29). The plot shows a mountainous curve with a peak around *Q* = 0.1 Å^-1^, suggesting that the rGli123 protein is not in a random coil but has a folded structure. For an ideal sphere, the Kratky plot shows a hyperbolic curve with a single peak in the small-angle regions. However, in the present data, the plot shows a shoulder around *Q* = 0.02 Å^-1^, marked by an inverted black triangle in the figure (Fig. 6C), suggesting the existence of another hidden peak. The bimodal character of the Kratky plot suggests a dumbbell shape for the protein; the innermost shoulder reflects the globularity of the whole molecule, while the peak reflects that of each domain (30). According to the pair-distance distribution function, *P*(*r*), the maximum length (*D*_max_) of the rGli123 protein was approximately 31.6 nm (Fig. 6D). The 3D structure of the rGli123 protein was modeled from SAXS data using PRIMUSQT software (Fig. 6E) (31). The structural model was rod-shaped and featured several narrow parts, similar to an articulated robot. This structure was consistent with the images obtained using rotary-shadowing EM with low ionic intensity (Fig. 4B). In the cross-section plot [ln *I*(*Q*)*Q vs. Q*^2^], which tests the validity of the rod-like structure in the SAXS model (32), the slope of the plot showed high linearity (Fig. S5). The radius of gyration of cross-section *R_c_* was 1.14 nm, which was obtained by fitting the cross-section plot to the equation ln [*I*(*Q*)*Q*] = ln [*I*(0)*Q*] – *R*_c_^2^*Q*^2^/2 (33). The radius of a cylinder *R*_p_ (a short-axis radius) was 1.61 nm, as calculated using the equation 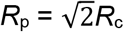 (33). The height of the cylinder *L* (twice the major-axis radius), determined from the equation *R*_g_^2^ = *R*_p_^2^/2 + *L*^2^/12 (33), was 28.2 nm. The *L* value obtained from the cross-section plot was consistent with the length (*D*_max_ = 31.6 nm) estimated from the *P*(*r*) function, indicating the high validity of the rod-like shape of the SAXS model. We also believe that the rod-like structure is likely to be a dimer, because the SAXS model has a molecular volume that corresponds to two molecules of a model predicted from the amino acid sequence using Robetta (Fig. 6F) (34). SAXS measurements were also performed using static cells at three different concentrations of rGli123 (Fig. S6). These results were consistent with those obtained from the SEC-SAXS measurements, and supported the absence of aggregation in the SAXS measurements.

**FIG. 6.**
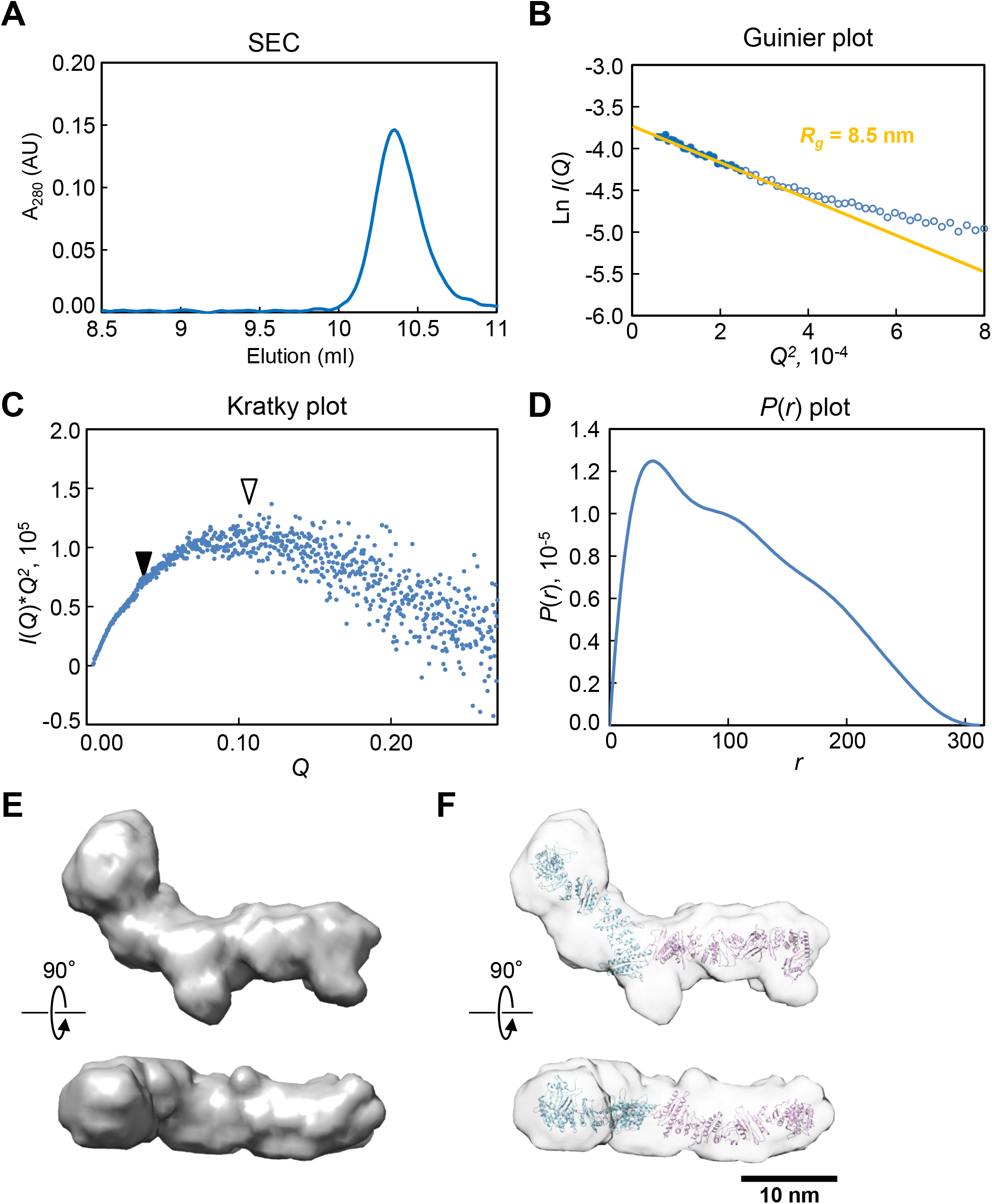
Rod-like structure of rGli123 protein, as predicted using SEC-SAXS. (A) SEC of rGli123 was carried out by measuring the UV absorbance at the wavelength of 280 nm, in the presence of 100 mM ammonium acetate. (B) Scattering intensities transformed as a Guinier plot [ln *I*(Q) *versus Q*^2^]. Data-points in a Guinier region (*R*_g_*Q* < 1.3), which are shown in the figure using filled circles, were then used to determine the *R*g value and zero-angle scattering intensity *I*(0) with Guinier approximation. (C) Scattering intensities transformed as a Kratky-plot [*I*(*Q*)*Q*^2^ *versus Q*]. White and black triangles indicate the peak and hidden peak (shoulder), respectively. (D) Pair-distance distribution function *P*(*r*) calculated from the scattering curve. (E) SAXS models of rGli123. (F) Overlay of SAXS models and the structures predicted from the amino acid sequence (1–1000 aa) using Robetta.

## DISCUSSION

### Surface gliding proteins

In this study, *gli123* and *gli42* genes were found in the genomes of *M. testudineum* and *M. agassizii* (Table S3), indicating that the two surface proteins of *M. mobile*, Gli123 and Gli42, are also essential for gliding, similar to Gli349 and Gli521 (20). The topologies of phylogenetic trees were common for all these proteins and 16S rDNA (20), suggesting that these mycoplasmas have acquired gliding motility in evolution, not through horizontal transfer (Fig. S1). In addition, the orthologs of *gli349* and *gli521* were positioned adjacent to the genomes of all gliding *mycoplasmas* (Fig. S2) (20), and the localization of Gli349 is dependent on that of Gli521 (13). These gliding proteins may form a complex immediately after synthesis and establish a gliding machinery in which binding and pulling are smoothly interlocked.

### Structure and conformational changes of Gli123

We isolated the Gli123 protein and found that its molecular structure changed from rod-like to globular depending on the ionic strength. This conformational change may occur in the flexible regions identified through limited proteolysis (Fig. S4). A pair of Gli123 molecules predicted from the amino acid sequence fit into both the rod-like structure obtained using SAXS and the globular structure obtained by means of single-particle analysis, suggesting that these structures of Gli123 are dimers (Figs. 5D and 6F).

The structure of the isolated molecule changes drastically with ionic strength; however, *M. mobile* can glide under different ionic strength environments, suggesting that this conformational change is unlikely to occur in the completed machinery (13). We observed that the purified rGli123 protein complemented the gliding of mutant cells lacking Gli123. Conformational changes in the Gli123 molecule may be related to this observation. These three gliding proteins have a localization hierarchy of Gli123-Gli521-Gli349, suggesting a physical interaction between these proteins at the cell surface (13). Gli123 might reach the target position on the cell surface as a flexible rod shape and bundle Gli521 and Gli349 molecules on the cell surface through a drastic conformational change, which is induced by the electrostatic fields made by the neighboring proteins, Gli349 and Gli521.

### Roles of the lipoprotein-17 domain

We found that a common repeat sequence of the lipoprotein-17 domain was shared by the Mvsps and gliding proteins. The relationship between the gliding machinery and Mvsps has been proposed based on the existence of a repeat comprising of approximately 100 amino acids (22, 24). In this study, we succeeded in determining the similarity between the repeat sequences. The existence of the lipoprotein-17 domain in Gli123, Gli349, and their orthologs may suggest that they originated from a common surface protein (Figs. 1A and 7A). The structure of the lipoprotein-17 domain in Gli123 and Mvsp predicted using AlphaFold2 featured three β-strands, which is consistent with previously reported structural features determined using nuclear magnetic resonance (NMR) and crystallography (Fig. S7A and S7B) (24). The predicted Mvsp structures suggest that an α-helix with a short loop connects the lipoprotein-17 domains to form a flexible chain (Fig. S7C). Although the detailed structures of Gli123 and Gli349 are still unknown, their flexibility may be derived from the structures common with the lipoprotein-17 domain and adjacent regions. Lipoprotein-17 domain may be useful for adjusting molecular length and flexibility (Fig. 7A). It may provide length variation adjustment to Mvsps and Gli349, which are involved in antigenic modulation and gliding leg, respectively. Flexible and long leg proteins may be advantageous for catching the host cell surface (17). In case of Gli123, it may provide length and flexibility for its role in clamping other surface gliding proteins on the machinery surface, where as many as 450 gliding units are packed (Fig. 7B). The gliding units fixed by Gli123 can be aligned in an ordered way for smooth and efficient gliding (13, 27).

**FIG. 7.**
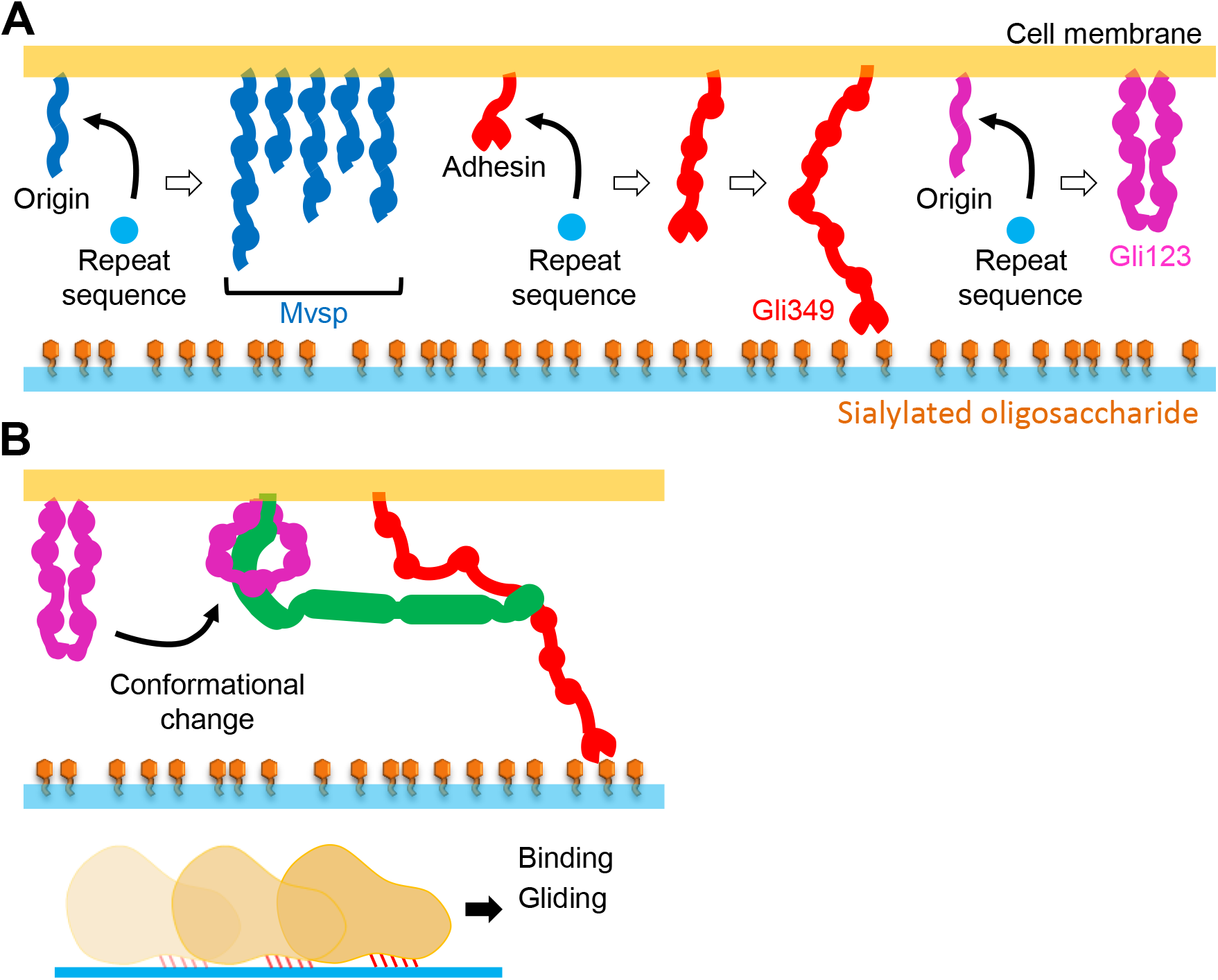
Role of lipoprotein-17 domain repeats and Gli123. (A) The repeat sequence is possibly used to adjust the length and flexibility of surface molecules that are involved in antigenic modulation (Mvsp) and gliding (Gli349 and Gli123). (B) The flexibility and molecular length of Gli123 allow for conformational changes in it, which facilitate its clamping to other gliding proteins at appropriate positions in the machinery.

## MATERIALS AND METHODS

### Sequence analyses

The genome sequences of *M. mobile* (163K), *M. pulmonis* (UAB CTIP), *M. agassizii* (PS6), and *M. testudineum* (ATCC700618) were obtained from the National Center for Biotechnology Information (NCBI) GenBank database (https://www.ncbi.nlm.nih.gov/genome/). A homology search against the NCBI non-redundant protein sequence databank was then performed using PSI-BLAST (https://blast.ncbi.nlm.nih.gov/Blast.cgi), with an expected threshold of 100. Phylogenetic trees of the proteins were constructed using neighbor-joining and maximum likelihood methods. The reliability of the topology of each phylogenetic tree was examined with 1000 bootstrap replicates, using the MEGAX package (https://www.megasoftware.net/). The secondary structure of Gli123 was predicted using XtalPred (https://xtalpred.godziklab.org/XtalPred-cgi/xtal.pl). Hidden Markov model scores were obtained from Pfam (http://pfam.xfam.org/family/PF04200) and UniProtKB (https://ftp.uniprot.org/pub/databases/uniprot/previous_releases/2021_03), based on the HMMER3 package (http://hmmer.org/).

### Strains and culture conditions

*M. mobile* 163K (ATCC 43663) was grown at 25°C in Aluotto medium, to an optical density at the wavelength of 600 nm (OD600) of around 0.08, as previously described (35). *The M. mobile* MvspI^-^ strain was isolated as follows. Mutants that did not react with monoclonal antibodies against MvspI were selected using colony blotting and Amido Black staining (36). The colonies were isolated and cultured in Aluotto liquid medium, following which SDS-PAGE analysis of the isolates was carried out, which confirmed the absence of MvspI protein. *E. coli* DH5α and BL21(DE3) strains were used for DNA recombination and protein expression, respectively.

### Isolation of Gli123 from *M. mobile* cells

The nGli123 protein was purified from the *M. mobile* MvspI^-^ strain cells. Cells stored at – 80°C were precultured and grown in 3 L of Aluotto liquid medium, dispensed into nine 300 cm^2^ tissue culture flasks, and statically cultured at 25°C until the OD_600_ reached 0.08. The cells were collected by means of centrifugation at 18,800 × *g* and 4°C, for 30 min, and washed twice with phosphate-buffered saline (PBS; 58 mM Na_2_HPO_4_, 17 mM NaH_2_PO_4_, and 68 mM NaCl). The precipitate was suspended in 30 mL of buffer A (20 mM Tris-HCl pH 8.0 and 0.1 mM phenylmethylsulfonyl fluoride) and sonicated for 2 min, at intervals of 5 s, using a sonicator (US-600T, Nippon Seiki Co., Niigata, Japan), on ice.

The cell lysate was combined with 30 mL of buffer A containing 2% Triton™ X-100 (v/v) and shaken on ice for 1 h, to solubilize the proteins. The cell lysate was separated by means of centrifugation at 32,300 × *g* and 4°C, for 30 min, following which the obtained supernatant was applied to an anion-exchange column (HiTrap™ Q HP 5 mL; Cytiva, Tokyo, Japan) equilibrated with buffer A. Proteins bound to the column were eluted with a NaCl concentration gradient of 0–500 mM, at 4°C. The eluted fraction containing Gli123 was concentrated to a volume of 10 mL and subjected to a SEC column (HiLoad® 26/600 Superdex® 200 prep grade, Cytiva) equilibrated with SEC buffer (20 mM Tris-HCl pH 8.0 and 150 mM NaCl), at 4°C. Thyroglobulin (669 kDa), ferritin (440 kDa), aldolase (158 kDa), and ovalbumin (43 kDa), with Stokes radii of 8.5, 6.1, 4.8, and 3.1 nm, respectively, were used as standards. Protein fractions were analyzed using SDS-PAGE with a 10% polyacrylamide gel, followed by CBB staining.

### Expression and purification of recombinant Gli123

The N-terminal sequence of Gli123 was determined using Edman degradation, as previously described (11, 12). According to the sequencing results, the DNA sequence encoding the amino acid region from the 19^th^ to 1128^th^ position was codon-optimized for expression in *E. coli* and inserted between the *Bam*HI and *Xho*I sites in the multicloning site of the pET21a plasmid (Novagen, Madison, WI). The recombinant protein, rGli123, was expressed in *E. coli* BL21(DE3) and induced using 1 mM isopropylthio-β-galactoside at 15°C, for 18 h. Following that, the cells were collected, suspended in His-buffer A (20 mM Tris-HCl pH 8.0 and 20 mM NaCl), and washed twice.

The cells were suspended in 60 mL His-buffer A with 60 μL of 100 mM phenylmethylsulfonyl fluoride and sonicated for 4 min, at intervals of 5 s, on ice, as described above. The lysate was centrifuged at 32,300 × *g* and 4°C, for 30 min, following which the supernatant was subjected to Ni^2+^-NTA affinity chromatography (HisTrap™ HP 5 mL, Cytiva). Proteins bound to the column were eluted with an imidazole concentration gradient of 0–250 mM, at 4°C. The fraction containing rGli123 was adjusted to 20% saturation of ammonium sulfate, kept on ice for 1 h, following which the obtained supernatant was collected after centrifugation at 32,300 × *g* and 4°C, for 30 min. The supernatant was subjected to hydrophobic interaction chromatography column (Phenyl Sepharose HP 1 mL, Cytiva) and eluted in a gradient of 20%–0% ammonium sulfate. The rGli123 protein was concentrated to 5 mL by means of centrifugation at 5,000 × *g* and 4°C, using a 50k Amicon tube (Amicon Ultra, Merck, Darmstadt, Germany), and subjected to SEC. The protein concentration was adjusted using a 50k Amicon tube, if necessary, for analyses.

### Optical analyses

The circular dichroism spectra of the proteins were measured in the range of 200–250 nm using a spectropolarimeter (J-805, Jasco International Co., Tokyo, Japan), at 25°C. A quartz cuvette with a 1-mm path length was filled with 1 mg/mL of rGli123 in SEC buffer. The conditions for the measurements have been previously described (37). The measured spectrum was analyzed using BeStSel (26), followed by estimation of the α-helix, β-strand, and turn contents (25). The secondary structure was predicted from the amino acid sequence using XtalPred (38). For light scattering, the rGli123 protein dialyzed against 50 mM ammonium acetate was placed into a 10-mm quartz cell, and the scattered light at the wavelength of 400 nm was measured at 25°C using a fluorescence meter (FP-6200, Jasco International Co.). The ammonium acetate concentration was adjusted by means of addition of 3 M ammonium acetate solution.

### Optical and electron microscopy

Rotary-shadowing and negative-staining EM were performed as previously described (10, 19, 24, 39). Protein images were analyzed using the image analysis software EMAN2.91 (https://blake.bcm.edu/emanwiki/EMAN2). The approximately 10,000 particle images were selected and classified into 50 groups, averaged, and then reconstructed into 3D structures from 3,202 particle images. For optical microscopy, *M. mobile* cells were inserted into a tunnel slide (20 μm) precoated with Aluotto medium, for 60 min. After 1 min, the floating cells were removed. The cells on the glass surface were observed using phase-contrast microscopy and analyzed using ImageJ v1.51w (http://rsb.info.nih.gov/ij/), as previously described (17, 20, 40).

### Assessment of binding of rGli123 to mycoplasma cells

Cultured *M. mobile* WT and m12 cells were washed twice and suspended in PBS. The suspended cells were kept in PBS containing 0.5 mg/mL of rGli123 protein for 3 h, following which they were collected by means of centrifugation at 12,000 × *g* and 25°C, for 5 min, suspended in PBS, sonicated, and analyzed using SDS-PAGE with CBB staining. Protein band intensities were analyzed using ImageJ v1.51w.

### SAXS measurements

SEC-SAXS measurements were performed at beamline 10C of the Photon Factory of the High Energy Accelerator Research Organization (KEK), Tsukuba, Japan. The rGli123 proteins were adjusted to a concentration of 5.33 mg/mL and subjected to a SEC column (Superdex® 200 Increase 3.2/300, Cytiva) equilibrated with 100 mM ammonium phosphate, at 4°C. The camera configuration has been previously described (37). SAXS measurements using a static cell were performed at three different protein concentrations. The measurements of the two-dimensional scattering images were circularly averaged and converted to a one-dimensional scattering curve using the SAngler software (41). The scattering curves immediately before the elution of the protein were used as those of the blank samples. The average scattering curve of the blanks was subtracted from the scattering curve of the protein solution to obtain the scattering profile *I*(Q) of the protein. *R*_g_ and *I*(0) were obtained based on Guinier approximation within the Guinier region (*R*_g_*Q* < 1.3). The pairwise distance distribution function *P*(*r*) was calculated using GNOM software (31). The *ab initio* structural model was simulated using the DAMMIF and DAMMIN softwares (31). Briefly, the outline of the 3D structure was calculated 10 times independently with DAMMIF, using dummy atoms. The structural models were averaged using DAMAVER, and the averaged structural models were calculated again using DAMMIF, to obtain the final structural model. The constructed model was displayed in UCSF Chimera 1.10 (42). In the cross-section plot (Fig. S5), the radius of gyration of the cross-section (*R*_c_) was obtained by applying a linear approximation to a plot of the linear region in the range after *Q* = 1/*R_g_* (33). The radius of the short axis (*R_p_*) was calculated using the equation 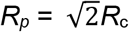 (33). The height of the cylinder (*L*) was calculated by substituting the values of *R_g_* and *R_p_* in *R_g_*^2^ = *R_p_*^2^/2 + *L*^2^/12 (33).

### Limited proteolysis

The rGli123 protein in SEC-buffer, at the concentration of 1 mg/mL, was treated with 0.001 mg/mL trypsin, at 25°C, for various reaction times, following which the reaction was stopped by means of heat treatment at 95°C for 5 min, in SDS-sample solution containing 5% glycerol, 0.025% Bromophenol Blue, 62.5 mM Tris-HCl pH 6.8, 2.5% SDS, and 5% β-mercaptoethanol. The protein digests were analyzed using SDS-PAGE with a 10% acrylamide gel and peptide mass fingerprinting, as described previously (19, 39).

## ACKNOWLEDGEMENTS

We thank Dr. Nobutaka Shimizu (KEK), Dr. Shinya Saijo (KEK), and the students of Arai Lab (Department of Life Sciences, The University of Tokyo) for their assistance with the SAXS measurements. We also thank Toshiaki Arata (Graduate School of Science, Osaka City University) and Ikuko Fujiwara (Department of Bioengineering, Nagaoka University of Technology) for their experimental advice in the light scattering analyses, and Ms. Aya Takamori (Graduate School of Science, Osaka City University) for her technical assistance with the MALDI-TOF MASS spectrometry. The SAXS measurements were performed with the approval of the Photon Factory Program Advisory Committee. This study was supported by Grants-in-Aid for Scientific Research on the Innovative Area Harmonized Supramolecular Motility Machinery and Its Diversity (MEXT KAKENHI grant numbers JP24117002, JP25117504, and JP15H01311), grants-in-aid for scientific research (B) and (A) (MEXT KAKENHI grant numbers JP24390107, JP17H01544, and JP19H02521), and JST CREST grant number JPMJCR19S5 to MM.

