## Supplementary material for "Structure and Function of Gli123 Involved in *Mycoplasma mobile* Gliding": Table S1 Table S2 Table S3 Table S4 FIG. S1 FIG. S2 FIG. S3 FIG. S4 FIG. S5 FIG. S6 FIG. S7

| Species | Strain | Protein | Annotation | Amino acid | Accession(NCBI) | Contig |
|---|---|---|---|---|---|---|
| *Mycoplasma mobile* | 163K | Gli123 | MMOB1020 | 1,128 | WP_011264622.1 | |
| | | Gli349 | MMOB1030 | 3.183 | WP_011264623.1 | |
| | | Gli521 | MMOB1040 | 4,727 | WP_011264624.1 | |
| | | Gli42 | MMOB1050 | 356 | WP_011264625.1 | |
| *Mycoplasma pulmonis* | UAB CTIP | Gli123 | MYPU_2160 | 1,128 | CAC13389.1 | |
| | | Gli349 | MYPU_2110 | 3,216 | CAC13384.1 | |
| | | Gli521 | MYPU_2120-2140 | 4,274 | CAC13385-13387 | |
| | | Gli42 | MYPU_2170 | 368 | CAC13390.1 | |
| *Mycoplasma agassizii* | PS6 | Gli123 | CFJ60_02770 | 1,188 | WP_084232031.1 | NQMN01000002 |
| | | Gli349 | CFJ60_05070 | 3,308 | WP_084232749.1 | NQMN01000003 |
| | | Gli521 | CFJ60_05075 | 4,582 | WP_084232748.1 | NQMN01000003 |
| | | Gli42 | CFJ60_02775 | 371 | WP_084232030.1 | NQMN01000002 |
| *Mycoplasma testudineum* | ATCC 700618 | Gli123 | EI74-0698 | 1,231 | WP_094254852.1 | SNWN01000014 |
| | | Gli349 | EI74_0476 | 3,190 | WP_094254640.1 | SNWN01000011 |
| | | Gli521 | EI74_0475 | 4,508 | WP_094254639.1 | SNWN01000011 |
| | | Gli42 | EI74-0697 | 403 | WP_094254851.1 | SNWN01000014 |

**Table S1**

Orthologs of *Mycoplasma mobile* genes coding surface gliding proteins.

| | sequences | regions |
|---|---|---|
| **Bacteria / Tenericutes / Mollicutes / Mycoplasmataceae (32) /** | **221** | **852** |
| *Mycoplasma pulmonis* (strain UAB CTIP) | 22 | 71 |
| *Mycoplasma hyorhinis* SK76 | 1 | 1 |
| *Mycoplasma bovigenitalium* 51080 | 1 | 1 |
| *Mycoplasma columbinum* SF7 | 5 | 43 |
| *Mycoplasma gallinarum* | 5 | 34 |
| *Ureaplasma parvum* serovar 3 (strain ATCC 700970) | 2 | 3 |
| *Mycoplasma mobile* (strain ATCC 43663 / 163K / NCTC 11711) | 22 | 85 |
| *Mycoplasma arthritidis* (strain 158L3-1) | 4 | 21 |
| *Mycoplasma orale* | 3 | 9 |
| *Mycoplasma bovoculi* M165/69 | 1 | 2 |
| *Mycoplasma phocidae* | 8 | 9 |
| *Mycoplasma meleagridis* ATCC 25294 | 6 | 15 |
| *Mycoplasma pullorum* | 5 | 32 |
| *Mycoplasma iowae* DK-CPA | 15 | 57 |
| *Mycoplasma hominis* (strain ATCC 23114 / NBRC 14850 / NCTC 10111 / PG21) | 1 | 8 |
| *Mycoplasma columborale* | 4 | 26 |
| *Mycoplasma columbinasale* | 23 | 103 |
| *Mycoplasma glycophilum* | 4 | 29 |
| *Mycoplasma gallinaceum* | 8 | 57 |
| *Mycoplasma cloacale* | 1 | 2 |
| *Mycoplasma gallopavonis* | 2 | 9 |
| *Mycoplasma alligatoris* A21JP2 | 3 | 5 |
| *Mycoplasma penetrans* (strain HF-2) | 19 | 72 |
| *Mycoplasma fermentans* (strain ATCC 19989 / NBRC 14854 / NCTC 10117 / PG18) | 2 | 5 |
| *Mycoplasma hyosynoviae* | 13 | 34 |
| *Mycoplasma subdolum* | 3 | 4 |
| *Mycoplasma agassizii* | 24 | 75 |
| *Mycoplasma crocodyli* (strain ATCC 51981 / MP145) | 3 | 12 |
| *Mycoplasma caviae* | 3 | 6 |
| *Mycoplasma maculosum* | 3 | 5 |
| *Mycoplasma phocicerebrale* | 4 | 11 |
| *Mycoplasma testudineum* | 1 | 6 |

**Table S2**
Lipoprotein-17 domain in *Mycoplasma*. The number of protein sequences containing the lipoprotein-17 domain (sequences) and the total number of these domains in these proteins (regions) of mycoplasma are shown based on analyses using Pfam version 33.1.

| Protein | sequence | | | Score (bits) | | E-value | |
|---|---|---|---|---|---|---|---|
| | Start | End | Length | Sequence | Domain | Sequence | Domain |
| MMOB1020 | 483 | 575 | 93 | 43.4 | 26.4 | 1.00E-07 | 2.00E-02 |
| ( *M. mobile* ) | 690 | 792 | 103 | | 13.8 | | 1.70E+02 |
| | 920 | 1016 | 97 | | 24.6 | | 7.50E-02 |
| | 1026 | 1127 | 102 | | 21.7 | | 6.20E-01 |
| MYPU2160 | 265 | 357 | 93 | 91.5 | 32 | 1.00E-22 | 3.70E-04 |
| ( *M. pulmonis* ) | 645 | 727 | 83 | | 24.5 | | 8.00E-02 |
| | 738 | 817 | 80 | | 23.7 | | 1.40E-01 |
| | 940 | 1030 | 91 | | 22.8 | | 2.70E-01 |
| | 1045 | 1127 | 83 | | 33.1 | | 1.60E-04 |
| CJF60-02770 | 252 | 351 | 100 | 59.4 | 13 | 3.80E-12 | 1.10E+03 |
| ( *M. agassizii* ) | 513 | 628 | 116 | | 15 | | 2.70E+02 |
| | 656 | 739 | 84 | | 11.3 | | 3.80E+03 |
| | 750 | 825 | 76 | | 16.6 | | 8.70E+01 |
| | 990 | 1081 | 92 | | 16.6 | | 8.50E+01 |
| | 1090 | 1187 | 98 | | 16.3 | | 1.10E+02 |
| EI74-0698 | 158 | 243 | 86 | 56.6 | 13.6 | 8.00E-12 | 2.00E+02 |
| ( *M. testudineum* ) | 258 | 364 | 107 | | 17 | | 1.80E+01 |
| | 678 | 771 | 94 | | 17 | | 1.80E+01 |
| | 775 | 865 | 91 | | 20.2 | | 1.80E+00 |
| | 1131 | 1230 | 100 | | 18.5 | | 6.00E+00 |
| MMOB1030 | 173 | 265 | 93 | 62.3 | 26 | 1.30E-13 | 2.80E-02 |
| ( *M. mobile* ) | 454 | 551 | 98 | | 27.2 | | 1.20E-02 |
| | 554 | 665 | 112 | | 19.9 | | 2.20E+00 |
| | 670 | 767 | 98 | | 24.4 | | 8.70E-02 |
| | 2153 | 2241 | 89 | | 27.2 | | 1.20E-02 |
| | 2460 | 2556 | 97 | | 20.9 | | 1.10E+00 |
| MYPU2110 | 49 | 148 | 100 | 70.7 | 23.7 | 3.20E-16 | 1.40E-01 |
| ( *M. pulmonis* ) | 445 | 542 | 98 | | 20.7 | | 1.30E+00 |
| | 551 | 644 | 94 | | 19.3 | | 3.50E+00 |
| | 658 | 747 | 90 | | 31.7 | | 4.60E-04 |
| | 1093 | 1181 | 89 | | 11.7 | | 8.00E+02 |
| | 1698 | 1785 | 88 | | 19.1 | | 4.00E+00 |
| | 2109 | 2197 | 89 | | 33.7 | | 1.10E-04 |
| | 2798 | 2887 | 90 | | 21.1 | | 9.30E-01 |
| CJF60-05070 | 352 | 440 | 89 | 48.1 | 12.6 | 1.30E-08 | 1.50E+03 |
| ( *M. agassizii* ) | 449 | 550 | 102 | | 12.9 | | 1.20E+03 |
| | 560 | 650 | 91 | | 23.6 | | 5.70E-01 |
| | 666 | 761 | 96 | | 18.4 | | 2.40E+01 |
| | 775 | 872 | 98 | | 11.9 | | 2.60E+03 |
| | 1112 | 1209 | 98 | | 17.8 | | 3.60E+01 |
| | 2158 | 2254 | 97 | | 30.7 | | 3.40E-03 |
| | 2842 | 2935 | 94 | | 17.7 | | 3.90E+01 |
| EI74-0476 | 562 | 650 | 89 | 33.9 | 20.3 | 9.60E-05 | 1.70E+00 |
| ( *M. testudineum* ) | 792 | 878 | 87 | | 11.3 | | 1.00E+03 |
| | 1119 | 1199 | 81 | | 13.1 | | 3.00E+02 |
| | 2126 | 2223 | 98 | | 24.8 | | 6.60E-02 |
| | 2788 | 2879 | 92 | | 22.3 | | 4.10E-01 |
| Mvsp A | 82 | 168 | 87 | 147.2 | 45.3 | 4.30E-40 | 2.60E-08 |
| | 173 | 294 | 122 | | 47.3 | | 6.00E-09 |
| | 304 | 393 | 90 | | 40.6 | | 7.60E-07 |
| | 403 | 490 | 88 | | 27.7 | | 8.00E-03 |
| Mvsp B | 46 | 128 | 83 | 181.7 | 54.6 | 7.10E-51 | 3.30E-11 |
| | 138 | 220 | 83 | | 59.4 | | 1.10E-12 |
| | 229 | 311 | 83 | | 58.4 | | 2.10E-12 |
| | 316 | 401 | 86 | | 39.7 | | 1.50E-06 |
| Mvsp C | 42 | 124 | 83 | 110.9 | 49 | 8.70E-29 | 1.90E-09 |
| | 131 | 217 | 87 | | 41.8 | | 3.30E-07 |
| | 227 | 304 | 78 | | 35.6 | | 2.80E-05 |

| Protein | sequence | | | Score (bits) | | E-value | |
|---|---|---|---|---|---|---|---|
| | Start | End | Length | Sequence | Domain | Sequence | Domain |
| Mvsp D | 49 | 131 | 83 | 146.2 | 44.8 | 8.40E-40 | 3.70E-08 |
| | 142 | 226 | 85 | | 37.5 | | 7.20E-06 |
| | 233 | 322 | 90 | | 48.6 | | 2.50E-09 |
| | 327 | 410 | 84 | | 47.3 | | 6.40E-09 |
| Mvsp E | 43 | 126 | 84 | 255.5 | 48.6 | 6.70E-74 | 2.50E-09 |
| | 139 | 224 | 86 | | 51 | | 4.40E-10 |
| | 236 | 322 | 87 | | 50.9 | | 4.50E-10 |
| | 338 | 420 | 83 | | 47.8 | | 4.40E-09 |
| | 431 | 518 | 88 | | 54.4 | | 3.70E-11 |
| | 523 | 607 | 85 | | 37.8 | | 5.60E-06 |
| Mvsp F | 43 | 126 | 84 | 247.7 | 46.5 | 1.90E-71 | 1.10E-08 |
| | 139 | 224 | 86 | | 51.1 | | 4.20E-10 |
| | 236 | 322 | 87 | | 50.9 | | 4.50E-10 |
| | 338 | 420 | 83 | | 47.6 | | 5.10E-09 |
| | 431 | 518 | 88 | | 57 | | 6.00E-12 |
| | 523 | 607 | 85 | | 38.9 | | 2.70E-06 |
| Mvsp H | 37 | 128 | 92 | 146.4 | 51.6 | 7.60E-40 | 2.90E-10 |
| | 134 | 218 | 85 | | 51.3 | | 3.50E-10 |
| | 229 | 318 | 90 | | 54 | | 5.00E-11 |
| Mvsp I | 38 | 129 | 92 | 623.6 | 50.8 | 9.20E-189 | 5.10E-10 |
| | 137 | 219 | 83 | | 54.2 | | 4.50E-11 |
| | 229 | 329 | 101 | | 49.8 | | 1.00E-09 |
| | 345 | 425 | 81 | | 44.9 | | 3.50E-08 |
| | 433 | 515 | 83 | | 43.5 | | 9.50E-08 |
| | 523 | 609 | 87 | | 45.3 | | 2.60E-08 |
| | 623 | 706 | 84 | | 43.6 | | 8.70E-08 |
| | 718 | 801 | 84 | | 41.5 | | 4.00E-07 |
| | 809 | 892 | 84 | | 41.9 | | 3.10E-07 |
| | 900 | 982 | 83 | | 64.6 | | 2.50E-14 |
| | 994 | 1076 | 83 | | 62.2 | | 1.40E-13 |
| | 1084 | 1166 | 83 | | 61.1 | | 3.10E-13 |
| | 1173 | 1252 | 80 | | 37.9 | | 5.50E-06 |
| | 1260 | 1338 | 79 | | 26.4 | | 2.00E-02 |
| | 1353 | 1436 | 84 | | 31.3 | | 6.10E-04 |
| | 1687 | 1766 | 80 | | 31.9 | | 4.00E-04 |
| Mvsp J | 33 | 121 | 89 | 83.7 | 43.8 | 2.70E-20 | 7.90E-08 |
| | 135 | 225 | 91 | | 44.8 | | 3.80E-08 |
| Mvsp K | 38 | 130 | 93 | 143.1 | 41.1 | 7.90E-39 | 5.30E-07 |
| | 143 | 229 | 87 | | 44.3 | | 5.30E-08 |
| | 241 | 327 | 87 | | 46 | | 1.60E-08 |
| | 341 | 424 | 84 | | 39.9 | | 1.20E-06 |
| Mvsp L | 37 | 124 | 88 | 120.2 | 44 | 1.10E-31 | 6.50E-08 |
| | 132 | 225 | 94 | | 17.2 | | 1.60E+01 |
| | 229 | 315 | 87 | | 34.6 | | 5.80E-05 |
| | 322 | 408 | 87 | | 39.9 | | 1.30E-06 |
| | 414 | 500 | 87 | | 31.6 | | 4.80E-04 |
| Mvsp M | 57 | 139 | 83 | 83.3 | 58.9 | 3.50E-20 | 1.50E-12 |
| | 144 | 239 | 96 | | 28.6 | | 4.30E-03 |
| Mvsp N | 70 | 152 | 83 | 134.2 | 59 | 4.80E-36 | 1.40E-12 |
| | 170 | 252 | 83 | | 55.8 | | 1.30E-11 |
| | 273 | 355 | 83 | | 30.1 | | 1.50E-03 |
| Mvsp O | 52 | 154 | 103 | 194.9 | 58.9 | 5.60E-55 | 1.50E-12 |
| | 173 | 254 | 82 | | 71.3 | | 2.10E-16 |
| | 273 | 355 | 83 | | 55.9 | | 1.30E-11 |
| | 376 | 458 | 83 | | 25.5 | | 4.00E-02 |
| Mvsp P | 30 | 139 | 110 | 56.5 | 56 | 8.20E-12 | 1.20E-11 |

**Table S3**

Hidden Markov model (HMM) scores for the lipoprotein-17 domains in Mvsps as well as the Gli123 and Gli349 orthologs, obtained using Pfam. The "sequence score" is the score calculated from all domains in the sequence, while the "domain score" is the score for each domain. The threshold values for the sequence and domain scores were 22.4 and 11.3, respectively.

| gene_product | gene_ID | Codon WT <-> mvspI- | Amino acid WT<->mvspI- | mutate_type | gene_start | gene_end | strand |
|---|---|---|---|---|---|---|---|
| COF family HAD hydrolase protein | MMOB1410 | GAA <-> AAA | E <-> K | nonsyn | 197320 | 198141 | - |
| expressed protein | MMOB0180 | TTT <-> TCT | F <-> S | nonsyn | 21463 | 21813 | - |
| conserved expressed protein | MMOB1700 | AGT <-> ATT | S <-> I | nonsyn | 232701 | 233795 | - |
| unspecified sugar ABC transporter binding protein | MMOB0360 | CCT <-> CTT | P <-> L | nonsyn | 51115 | 52590 | + |
| variable surface protein mvspM | MMOB6070 | ATT <-> ATC | I <-> I | syn | 746365 | 747093 | - |
| variable surface protein mvspN | MMOB6080 | ACT <-> ATT | T <-> I | nonsyn | 747389 | 748456 | - |
| variable surface protein mvspN | MMOB6080 | TTG <-> TTA | L <-> L | syn | 747389 | 748456 | - |
| variable surface protein mvspN | MMOB6080 | AAT <-> AAC | N <-> N | syn | 747389 | 748456 | - |
| variable surface protein mvspN | MMOB6080 | CTT <-> TTT | L <-> F | nonsyn | 747389 | 748456 | - |

| gene_product | gene_ID | Codon WT <-> mvspI- | deletion_site | mutate_type | gene_start | gene_end | strand |
|---|---|---|---|---|---|---|---|
| variable surface protein mvspI | MMOB3340 | AAT <-> A—T | 423447 | Deletion | 417670 | 423678 | - |

**Table S4**

Mutation points in the MvspI-deficient mutant. Nine single nucleotide polymorphisms and one deletion mutation were found in the MvspI$^-$ strain. The *mvspI* gene underwent a frameshift mutation, in which the 98$^{th}$ codon was replaced by a stop codon.

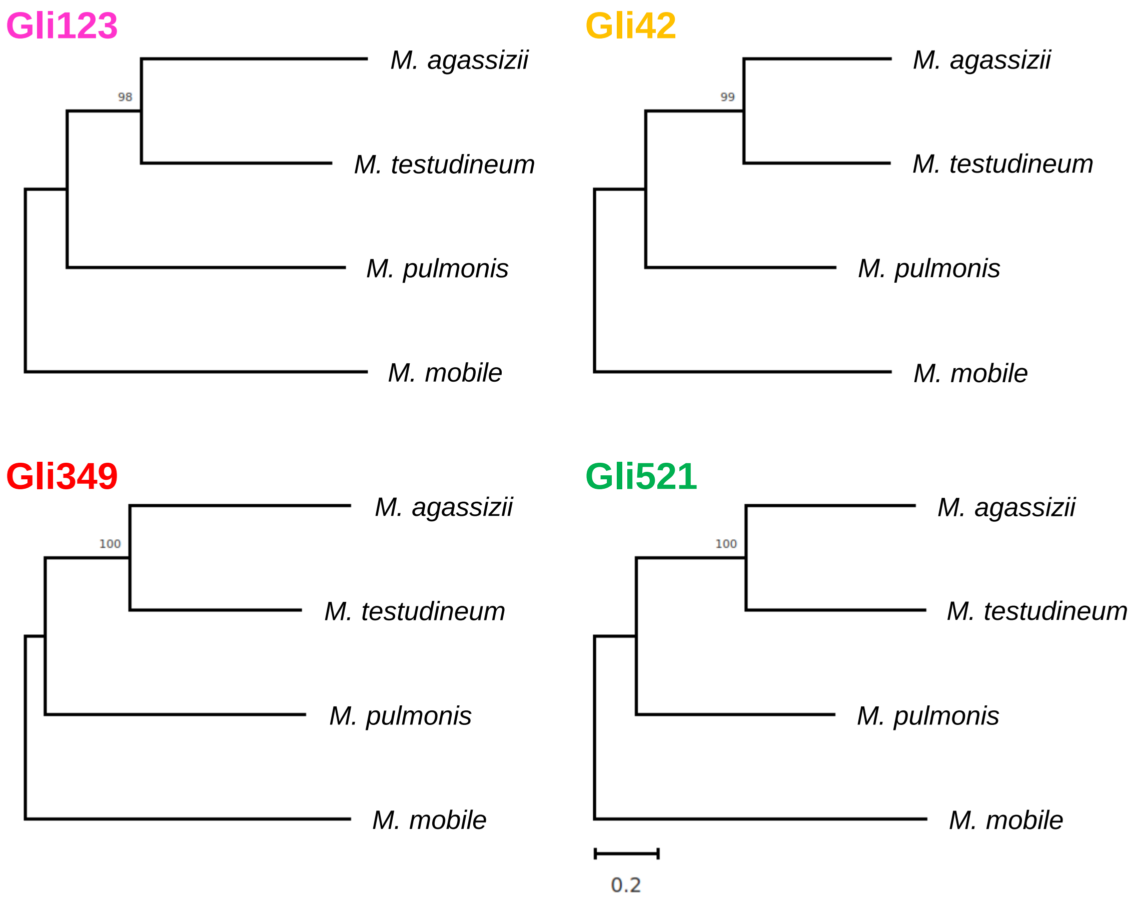


**FIG. S1** Phylogenetic trees of the surface proteins that constitute the gliding machinery. The phylogenetic trees were inferred using the maximum likelihood method and JTT matrix-based model, with the MEGAX package. The percentage of trees in which the associated taxa were clustered together is shown next to the branches. The trees have been drawn to scale, and the branch lengths indicate the number of substitutions per site.

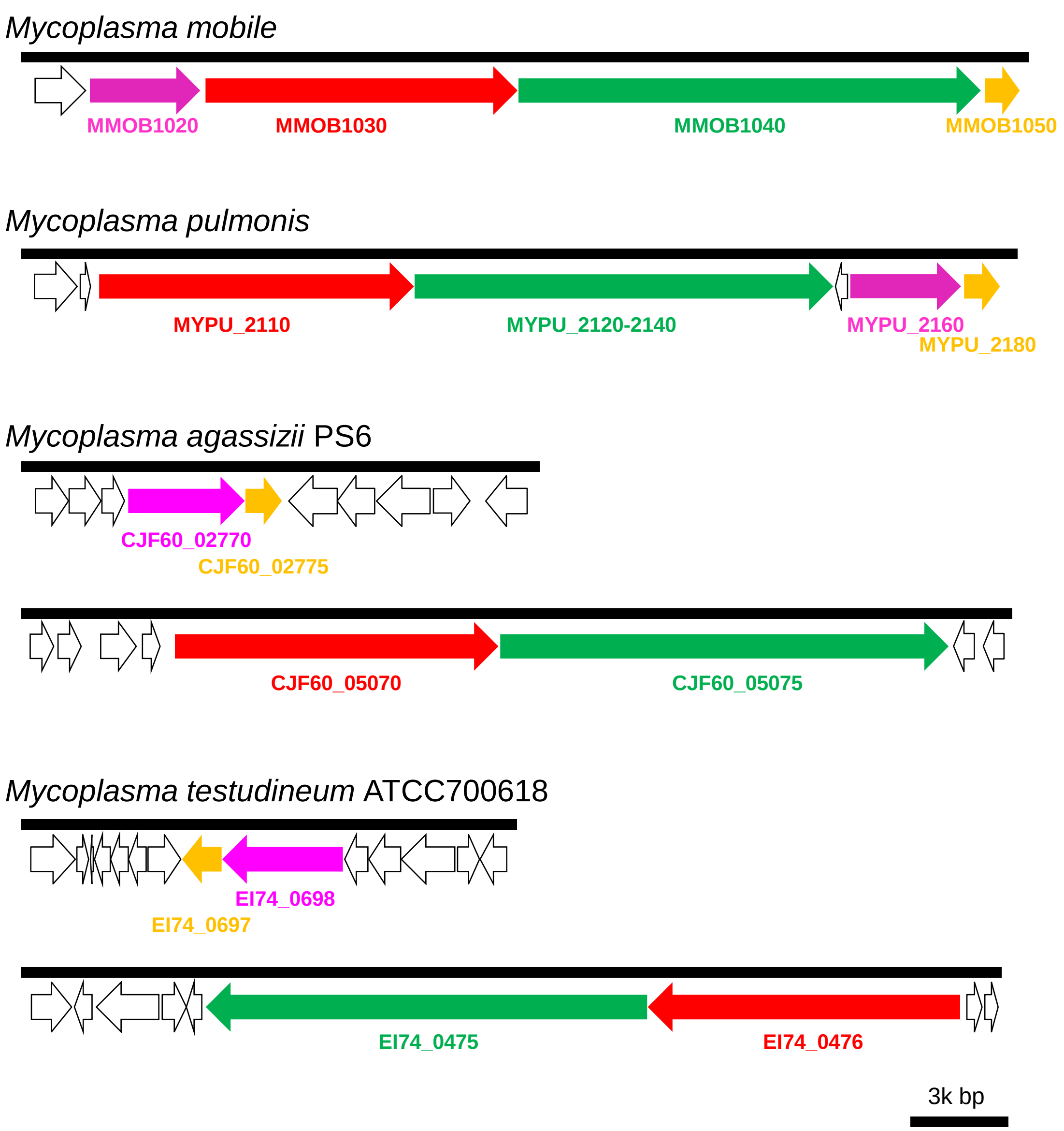


**FIG. S2** Gene loci coding putative surface proteins for gliding in the *M. mobile*, *M. pulmonis*, *M. agassizii*, and *M. testudineum* genomes. The magenta, red, green, and orange boxes show orthologs of *gli123*, *gli349*, *gli521*, and *gli42*, respectively.

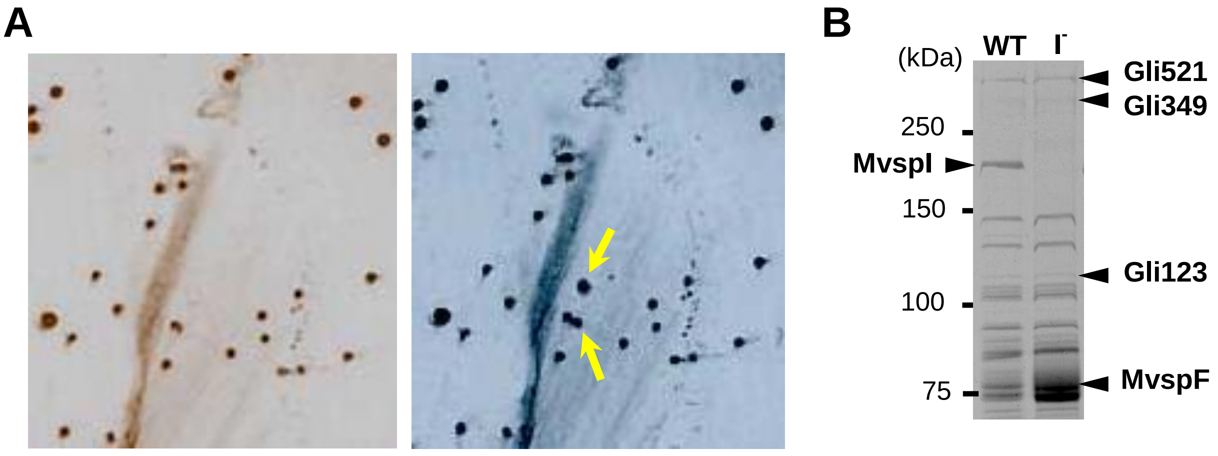


**C**

```
226 ACAAAAAATAATGCTACCACTATTAATGGTGTTGCCTTTTCTTATGCTCTTTCTGAAAAAACTCCTATAAATGCT   300 WT
     T  K  N  N  A  T  T  I  N  G  V  A  F  S  Y  A  L  S  E  K  T  P  I  N  A
226 ACAAAAA–TAATGCTACCACTATTAATGGTGTTGCCTTTTCTTATGCTCTTTCTGAAAAAACTCCTATAAATGCTG  300 MvspI- mutant
     T  K  I   M  L  P  L  L  M  V  L  P  F  L  M  L  F  L  K  K  L  L  *  M  L
```

**FIG. S3** Isolation of MvspI-defective mutant. (A) Colonies blotted to a sheet were tested for the presence of MvspI using a monoclonal antibody (left) and stained using Amido Black (right). MvspI-lacking colonies are marked using yellow arrows. (B) The protein profile of an isolated mutant. The whole cell lysate was subjected to SDS-PAGE with a 6% acrylamide gel. (C) Mutations in the gene encoding MvspI. The red hyphen indicates the point of mutation, while asterisk indicates the stop codon.

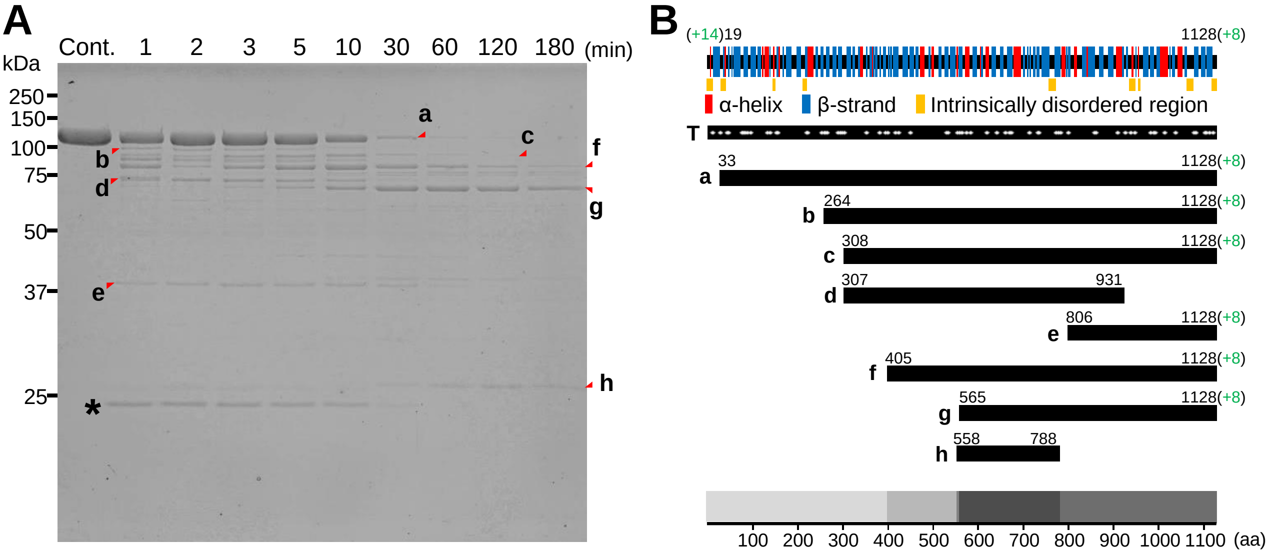


**FIG. S4** Limited proteolysis of rGli123. (A) Protein profiles of rGli123 digested various times with trypsin and analyzed by means of SDS-PAGE with a 10% acrylamide gel. The digested peptide fragments were identified as a–h. The trypsin band is marked using an asterisk. (B) The positions of the secondary structures predicted using XtalPred are given at the top. The trypsin-cleavable sites are indicated using white dots in (T). The degradation products a–h were assigned on the sequence based on the results of the peptide mass fingerprint analyses. The sequence regions of peptide bands (a–h) were assigned using both the evident residue numbers from PMF and the estimated band sizes, as previously described (39). The overlapping sequences of the degradation products (f, g, and h) suggest the rigidity of the molecule, as presented by a dark shade in the bottom schematic.

A

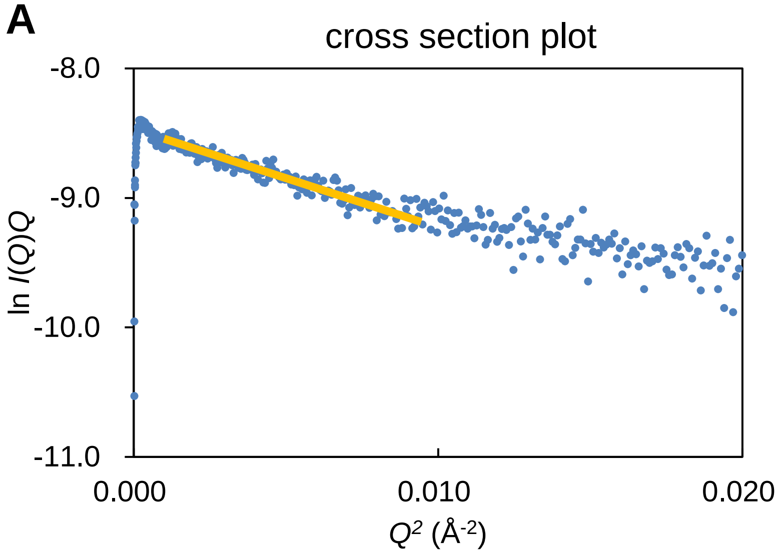


**FIG. S5** Validity of the rod shape of the Gli123 SAXS model. Scattering intensities transformed as a cross-section plot (ln *I*(*Q*)*Q* versus $Q^2$), where *I*(*Q*) and *Q* indicate scattering intensity and scattering vector, respectively.

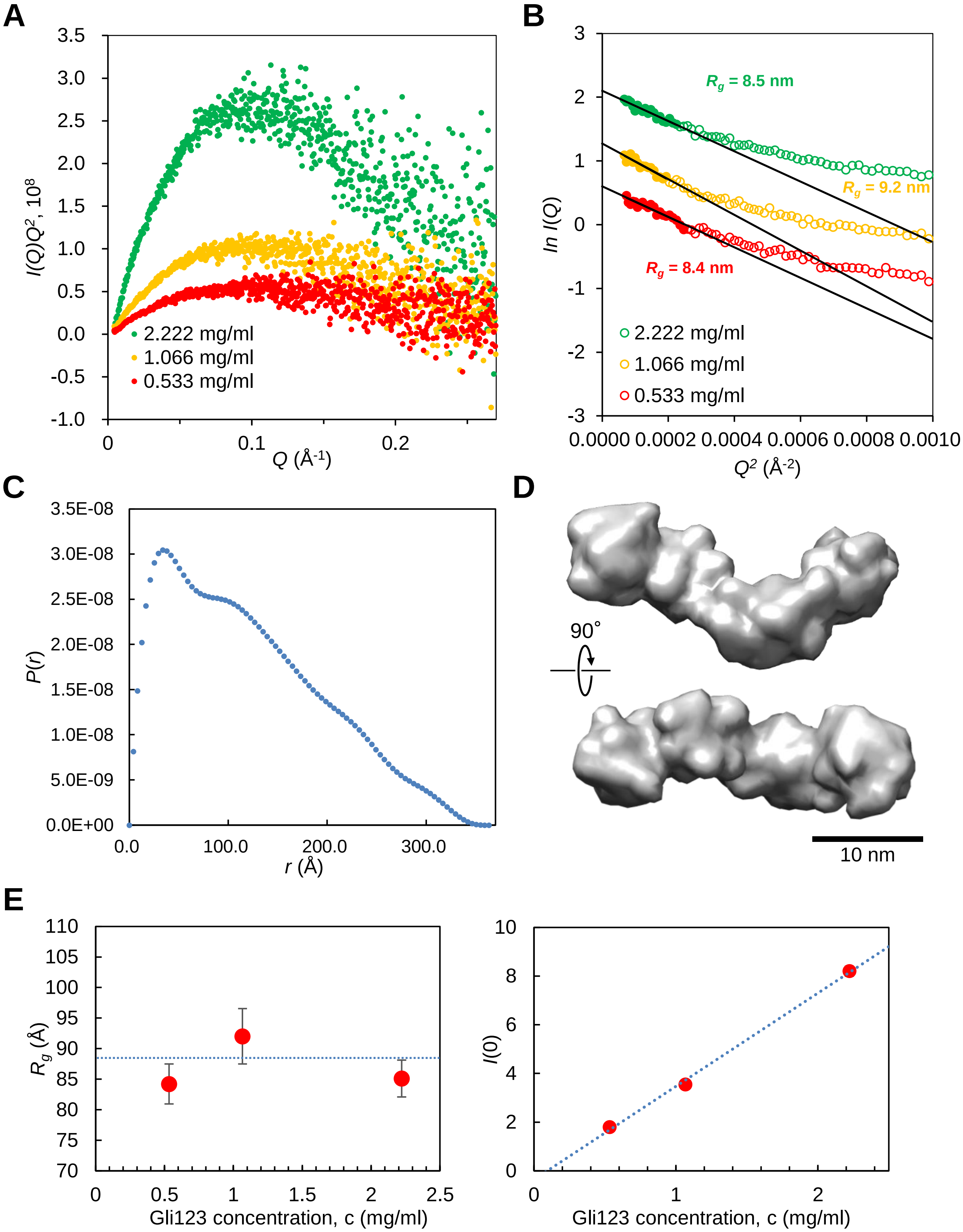


**FIG. S6** Small angle X-ray scattering of rGli123. SAXS measurements were performed using a static cell, at rGli123 concentrations of 0.533, 1.066, and 2.222 mg/mL. (A) Kratky plot ($I(Q)Q^2$ *versus* $Q$) and (B) Guinier plot (ln $I(Q)$ *versus* $Q^2$) of the scattering curve. Approximate lines were determined within the low $Q$-range of data limited by $R_gQ < 1.3$, indicated using filled color circles. The slopes have been calculated using the formula $-R_g^2/3$. The extrapolated intensity at zero angle is $I(0)$. (C) Pair-distance distribution function $P(r)$. (D) SAXS models of rGli123. (E) Concentration dependence of $R_g$ (left) and $I(0)$ (light) calculated from the Guinier plots. $I(0)$ is proportional to protein concentration, showing the absence of aggregation.

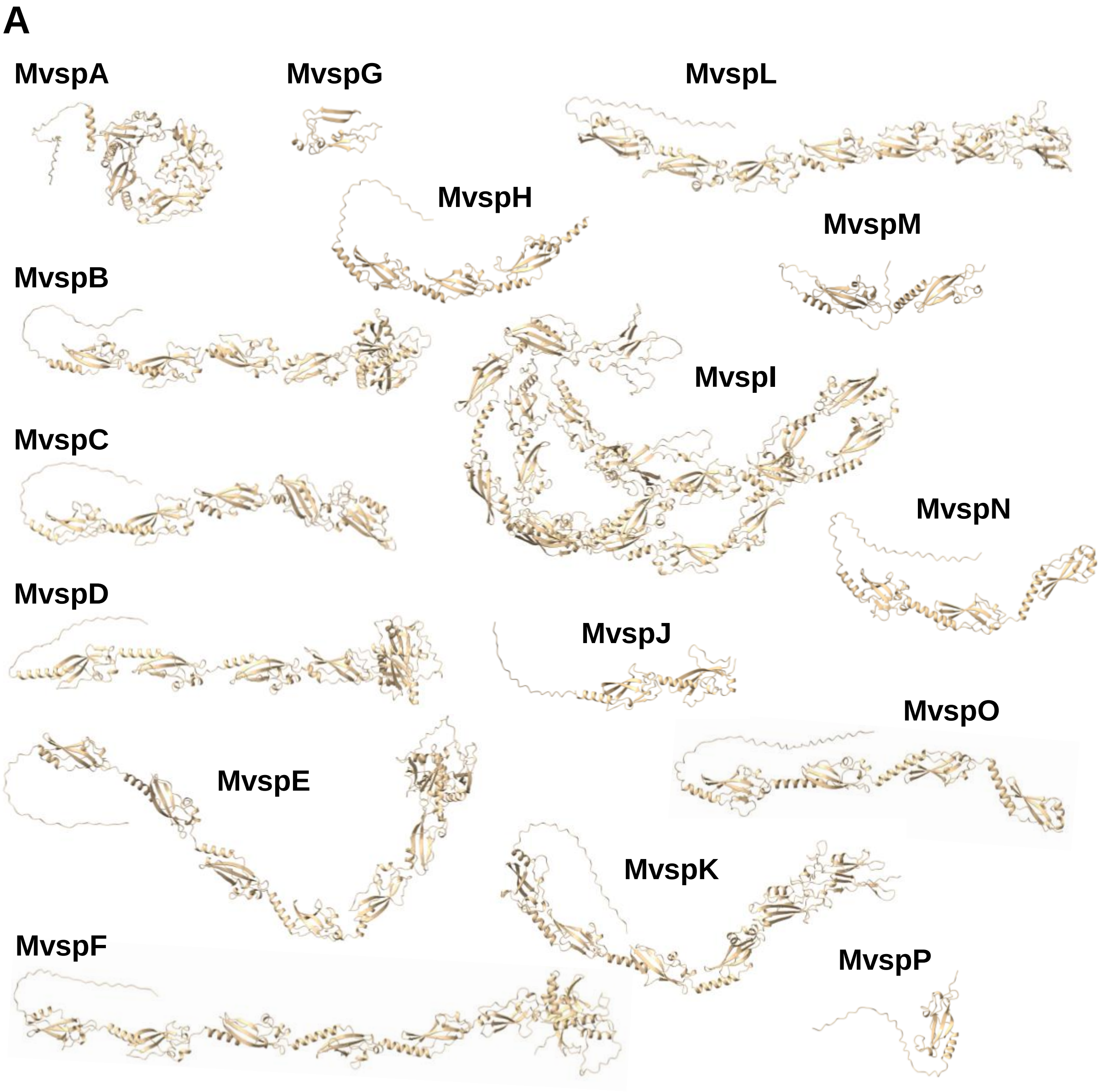


**FIG. S7** (A) Mvsp structures predicted using AlphaFold2.

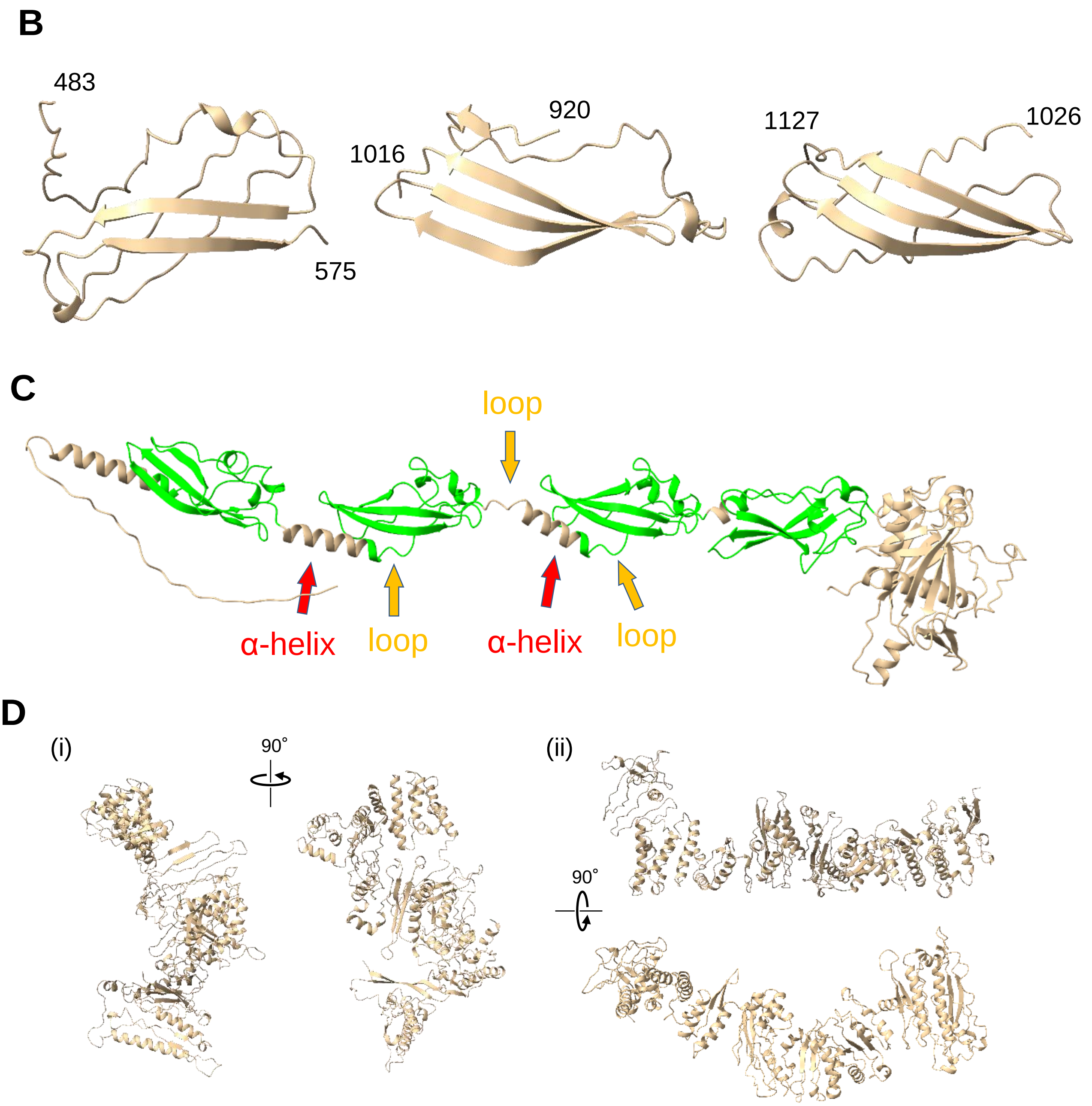


**FIG. S7** (B) Structure prediction of Gli123 lipoprotein-17 domain predicted using AlphaFold2. The predictions on the left, middle, and right are for the first, third, and fourth lipoprotein-17 domains of Gli123, respectively. (C) Structure of MvspD predicted using AlphaFold2. The lipoprotein-17 domains are colored green. The domains are connected by an alpha helix and a short loop. (D) Structure of Gli123 predicted using Robetta. (i) and (ii) show candidate predictions fitted to globular (Fig. 5) and rod-like (Fig. 6) structures, respectively.
